# The spliced leader RNA silencing (SLS) pathway in *Trypanosoma brucei* is induced by perturbations of endoplasmic reticulum, Golgi, or mitochondrial proteins factors and functional analysis of SLS inducing kinase, PK3

**DOI:** 10.1101/2021.09.07.459367

**Authors:** Uthman Okalang, Bar Mualem Bar-Ner, K. Shanmugha Rajan, Nehemya Friedman, Saurav Aryal, Katarina Egarmina, Ronen Hope, Netaly Khazanov, Hanoch Senderowitz, Assaf Alon, Deborah Fass, Shulamit Michaeli

## Abstract

In the parasite *Trypanosoma brucei*, the causative agent of human African sleeping sickness, all mRNAs are *trans*-spliced to generate a common 5’ exon derived from the spliced leader RNA (SL RNA). Perturbations of protein translocation across the endoplasmic reticulum (ER) induce the spliced leader RNA silencing (SLS) pathway. SLS activation is mediated by a serine-threonine kinase, PK3, which translocates from the cytosolic face of the ER to the nucleus, where it phosphorylates the TATA binding protein TRF4, leading to the shut-off of SL RNA transcription, followed by induction of programmed cell death. Here, we demonstrate that SLS is also induced by depletion of the essential ER resident chaperones BiP and calreticulin, ER oxidoreductin 1 (ERO1), and the Golgi-localized quiescin sulfhydryl oxidase (QSOX1). Most strikingly, silencing of *Rhomboid-like 1(TIMRHOM1)* involved in mitochondrial protein import, also induces SLS. The PK3 kinase, which integrates SLS signals, is modified by phosphorylation on multiple sites. To determine which of the phosphorylation events activate PK3, several individual mutations or their combination were generated. These mutations failed to completely eliminate the phosphorylation or translocation of the kinase to the nucleus. The structure of PK3 kinase and its ATP binding domain were therefore modeled. A conserved phenylalanine at position 771 was proposed to interact with ATP, and the PK3^F771L^ mutation completely eliminated phosphorylation under SLS, suggesting that the activation involves most if not all the phosphorylation sites. The study suggests that the SLS occurs broadly in response to failures in protein sorting, folding, or modification across multiple compartments.

## INTRODUCTION

*Trypanosoma brucei* is a digenetic parasite that cycles between the tsetse fly and a mammalian host. Trypanosomatids are known for their non-conventional gene expression mechanisms such as *trans-*splicing (1) and mitochondrial RNA editing (2). Trypanosomal mRNAs undergo *trans*-splicing whereby a small exon, the spliced leader (SL), provided by the small SL RNA, is donated to all pre-mRNAs by *trans*-splicing (1).

Conventional regulation of protein coding genes based on transcription from defined promoters is absent in trypanosomes, and regulation of gene expression is mostly post-transcriptional (3). It was consequently expected that these parasites would lack a typical unfolded protein response (UPR) mechanism, which in other eukaryotes involves transcriptional activation (4–6). Trypanosomes instead cope with ER stress, such as that induced by treatment with a reducing agent, by preferential stabilization of mRNAs that are essential for their ability to withstand such stress (7).

Under severe stress induced by silencing of mRNA encoding proteins involved in protein translocation into the ER lumen, such as the signal-recognition particle (SRP) receptor (*srα*), *sec61*, the translocation channel, and *sec63*, a factor involved in translocation to the ER, the spliced leader RNA silencing (SLS) pathway is induced (8, 9). SLS has two hallmarks: the reduced abundance of SL RNA, and an increased level of SNAP2, an SL RNA specific transcription factor that fails to bind to the SL RNA gene promoter and spreads throughout the nucleus (7, 9). We subsequently showed that many of the factors involved in the pre-initiation complex of SL RNA behave like SNAP2(10). SLS leads to programmed cell death (PCD) (7). Based on these data, we proposed that SLS triggers a unique death pathway similar to the classical caspase-mediated PCD observed in higher eukaryotes (4–6).

The mechanism underlying SLS was revealed by purifying the SL RNA transcription complex under SLS. It was found that the TATA-binding protein, TRF4, undergoes phosphorylation on serine 35 by serine-threonine-kinase, which was purified with the SL RNA transcription complex, and we termed it PK3. PK3 translocates from the face of the ER to the nucleus during SLS. PK3 is responsible for the activation of the PCD induced by SLS (10).

It is currently unknown if SLS is activated only by perturbations on the ER membrane or whether it is induced by depletion of factors involved in protein modification upon translocation into the ER. A number of conserved factors contribute to the three main modifications of proteins as they enter the ER: folding, disulfide bond formation, and glycosylation. The major ER chaperones that assist in protein folding are calnexin/calreticulin and BiP (immunoglobulin heavy chain binding protein) (11). Unlike calnexin/calreticulin, which monitor both N-linked glycans and unfolded regions on nascent polypeptide chains (12), BiP detects only the latter and is the major contributor to folding of non-glycosylated proteins (13). Trypanosomes like other eukaryotes possess the ER chaperones BiP and calreticulin (CRT) (14).

To accomplish disulfide bond formation, protein disulfide isomerase (PDI) oxidizes client proteins, and endoplasmic reticulum oxidoreductin 1 (ERO1) transfer electron from the reduced PDI to the terminal acceptor, which is usually molecular oxygen and that is subsequently reduced to H2O2. ERO1 function is essential for disulfide bond formation in yeast (15). As the oxidative activity of ERO1 is related to the production of H2O2 and burdens cells with potentially toxic ROS, deregulated ERO1 activity is likely to impair cell fitness (15).

Many proteins that are translocated to the ER undergo glycosylation, which is used both for its effect on protein stability and solubility, and as a code to monitor protein folding by ER quality control (ERQC). If a newly synthesized protein folds properly and passes the scrutiny of the ERQC machinery, it can be trafficked beyond the ER. Should the protein fail this inspection, it is targeted for proteasomal degradation in the cytosol via ER-associated degradation (ERAD) (16). In most eukaryotes, the glycan added to proteins in the ER is Glc(3) Man(9) GlcNAc(2), but in trypanosomes it is Glc1Man9GlcNAc2 (17). In the folding cycle, glucose is removed from this glycan by glycosidases. Trypanosomes encode for only a single glucosidase II (GLU2)(18). If the first round of folding fails, another cycle of folding is initiated by adding back the glucose via the enzyme UDP-glucose: glycoprotein glucosyltransferase (UGGT) (19). When misfolded proteins accumulate in the ER, and to avoid possible aggregation, another enzyme, α1,2-mannosidase, removes a mannose residue to produce Man8GlcNac2, which is then recognized by the protein ER degradation– enhancing alpha mannosidase (EDEM) (20), and the protein is tunneled to ERAD (21). In cases when the protein is properly folded, the protein moves out of the ER to the Golgi complex through the tubulovesicular membrane clusters of the ER-Golgi intermediate compartment (ERGIC). The ERGIC clusters are mobile transport complexes that deliver secretory cargo from ER-exit sites to the Golgi. The ERGIC also contributes to the concentration, folding, and quality control of newly synthesized proteins (22).

Trypanosomes cycle between insect and mammalian hosts. In the insect host, the primary surface protein is the procyclin EP, which covers the cell surface of the procyclic form (PCF) parasite, whereas in the bloodstream form (BSF) the main surface protein is the variant surface glycoprotein (VSG), which undergoes antigenic variation (23). In BSF, protein translocation across the ER is very active because of the massive production of VSG (24, 25). RNAi silencing of *bip*, *crt, glu2* and *uggt* mRNAs in BSF affected growth and resulted in a swollen ER (26).

A recent proteomics analysis of SLS identified a dramatic increase in the level of mitochondrial Rhomboid-like 1 (TIMRHOM1) (27). Recent studies suggest that this protein is involved in protein translocation to the mitochondria and may be a functional homologue of the mitochondrial pore, TIM23 (28). In eukaryotes, mitochondria are connected to the ER, and homeostasis is maintained between these two cellular domains. In yeast, this contact is mediated by the endoplasmic reticulum (ER)- mitochondrial encounter structure (ERMES) complex (29), and in mammals this contact involves the voltage-dependent anion selective channel protein (VDAC1), which interacts with ER Ca^2^+ channel IP3R (30). In trypanosomes, it is not known which proteins make the contact between these compartments.

In this study, we show that SLS is induced when perturbation in protein sorting or modification is elicited by depletion of factors located not only on the ER membrane but also inside the ER lumen. Silencing of *bip, crt, ero1 mRNAs* and a Golgi-localized oxidoreductase (*qsox1*) mRNA in PCF each elicited SLS, as demonstrated by reduction in the level of SL RNA, accumulation of TRF4 in the nucleus, and TRF4 phosphorylation. However, depletion of UGGT, GLU2, EDEM and ERGIC involved in protein trafficking did not elicit SLS, suggesting that not all the factors present in the ER are essential for protein homeostasis in PCF. In addition, we found that perturbation of proteins outside the ER can induce SLS, such as depletion of the mitochondrial TIMRHOM1. We further show that PK3, the kinase activated to initiate SLS, undergoes phosphorylation on multiple sites during SLS activation. The sites were mapped by mass spectrometry and verified by mutagenesis. Mutations introduced in S606, 607, 707, 708, and the combination of four mutations including S606, 608, 707, and 708 did not completely abolish the phosphorylation and translocation of PK3 to the nucleus. Modeling of the PK3 kinase domain based on the homologous mammalian PERK, allowed us to identify the ATP binding domain of PK3. Mutation of PK3^F771L^ in the ATP binding domain completely abolished the phosphorylation under SLS induction, further demonstrating that PK3 phosphorylation is essential for its translocation to the nucleus. Thus, this study provides evidence that SLS is activated under stress originating from inactivation of functions involved in protein sorting, folding, or modification, but not upon functional inactivation of all essential functions in the cell. SLS induction is not restricted to the ER, but encompasses other organelles of the secretory pathway and the mitochondria.

## MATERIALS AND METHODS

### Cell growth and transfection

Procyclic forms of *T. brucei* strain 29-13, which carries integrated genes for T7 polymerase and the tetracycline repressor (31), were grown in SDM-79 (32) supplemented with 10% fetal calf serum in the presence of 50 μg ml^-1^ hygromycin B and 15 μg ml^-1^ G418. Transfected cells were cloned by serial dilution to obtain a clonal population (31).

### Generation of transgenic parasites

Listed in Table S1 are the primers used to generate constructs for silencing the following genes: *bip* (Tb927.11.7460), *crt* (Tb927.4.5010), *glu2* (Tb927.10.13630), *uggt* (Tb927.3.4630), *ergic* (Tb927.11.4200), *edem* (Tb927.8.2910), *ero1* (Tb927.8.4890), *qsox1* (Tb927.6.1850), *timrhom1* (Tb927.9.8260). For most of the genes we generated stem-loop constructs, but T7 opposing constructs were also used (31). Stem-loop RNAi constructs were linearized by *EcorV* digestion and T7 opposing constructs were linearized with *NotI*. The PTP-tagged proteins were prepared with PCR products amplified using primers listed in Table S1, and cloned into the PTP vector (33).

### PK3 construct preparation and site-directed mutagenesis

The construct for tagging PK3 (Tb927.6.2980) with the C-terminal composite PTP tag was generated by cloning the PK3 gene fragment ∼1600 nt (amplified using the primers in Table S1) into the *ApaI* and *NotI* sites of pC-PTP-PURO (a kind gift from Prof. Laurie K. Read, University at Buffalo) (34), a derivative of pC-PTP-NEO in which the neomycin phosphotransferase coding region was replaced with that of puromycin N-acetyl-transferase (33). The resultant PK3-PTP vector was amplified in Dam^-/-^ bacteria, linearized with *BclI* (NEB), transfected, and cloned into cells carrying the *sec63* stem-loop RNAi construct as previously described (8). The construct for tagging PK3 with the C-terminal MYC tag was generated as described above using primers listed in Table S1 into the *HindIII* and *XbaI* sites of pNAT plasmid carrying MYC tag (a kind gift from Dr. Sam Alsford, LSHTM)(35). Site-directed mutagenesis was performed using the primers listed in Table S1. The template plasmid was digested with *DpnI* (NEB), and clones were isolated. Mutations in the plasmids and isolated procyclic clones were verified by Sanger sequencing.

### Northern analysis and primer extension

Total RNA was prepared with Trizol reagent, and 20 μg/lane was separated on a 1.2% agarose gel containing 2.2 M formaldehyde. Small RNAs were fractionated on a 6% (w/v) polyacrylamide gel containing 7 M urea. Primer extension was performed as previously described (36, 37). The extension products were analyzed on a 10% denaturing polyacrylamide gel. RNA probes were prepared by *in-vitro* transcription using α-^32^P-UTP. Primers used for *in-vitro* transcription are listed in Table S1.

### PK3 purification and mass-spectrometry analysis

Cells expressing the *SEC63* silencing construct and PK3 PTP- tagged protein (27), un- induced or induced for 2.5 days, were grown to a density of 2×10^7^cells ml^-1’^. The purification protocol used was essentially as described previously (33) from 10^10^ cells but included the addition of phosphatase inhibitors to the extract (10 mM NaF, 1 mM sodium orthovanadate and 50 mM beta-glycerophosphate). In addition, the lysis buffer was modified to contain 0.5% deoxycholate, 0.5% NP-40%, and 0.1% SDS to release the kinase from the membrane. After separation of purified proteins by denaturing 8% SDS-PAGE, protein bands were eluted from the gel and subjected to trypsin digestion. The resulting peptides were resolved by reverse-phase chromatography. Mass spectrometry was performed by an ion-trap mass spectrometer (Orbitrap XL, Thermo). To analyze phosphopeptides, the proteins were enriched on a TiO2 column. The mass spectrometry data were analyzed using the Discoverer software version 1.2 or 1.3 against the *T. brucei* TriTrypdb database (http://tritrypdb.org/tritrypdb/).

### Preparation of TRF4 and TIMRHOM1 antibodies and source of antibodies

The *trf4* gene and a portion of the *timrhom1* gene were amplified by PCR using primers listed in S-1. The amplified fragments were cloned into the pHIS vector (Novagen) and expressed in *E. coli* BL21 cells. Recombinant proteins were extracted and purified using the Bugbuster reagent (Novagen, Inc). To raise antibodies against TRF4 or TIMRHOM1, 400 µg of the proteins were emulsified with an equal volume of complete adjuvant (Difco). The emulsions were injected subcutaneously into two female New Zealand white rabbits. The first injection was followed by an additional two injections of 200 µg protein emulsified with an equal volume of incomplete adjuvant (Difco) at 2-week intervals. Serum was collected and examined for reactivity by immunofluorescence and western analysis.

GRASP, SEC24, and GRIP70 antibodies were kindly provided by Dr. Graham Warren, (Vienna University, Austria) (38) (39), QSOX1 antibody was prepared by the group of Prof. D. Fass (Weizmann Institute, Israel) (40), PTB1 and ZC3H41 antibodies were prepared in our lab (41)(42), HSP83 antibody made against the Leishmania protein was kindly provided by Dr. Dan Zilberstein (Technion, Israel), and VH+ppase antibody was kindly provided by Dr. Roberto Docampo (University of Georgia, USA). EP antibodies was purchased from Cedarlane, Canada. MYC (9E10) antibody was purchased from Santa-Cruz. The dilutions used are specified in the Figure legends.

### Western analysis

Whole-cell lysates (10^7^ cells) were fractionated by 10% SDS-PAGE, transferred to PROTRAN membranes (Whatman), and reacted with antibodies. The bound antibodies were detected with goat anti-rabbit immunoglobulin G coupled to horseradish peroxidase and were visualized by ECL (Amersham Biosciences).

### Immunofluorescence assay

Cells were washed with PBS, mounted on poly-L-lysine-coated slides, and fixed in 4% formaldehyde. Immunofluorescence was performed as described (9). Cells were visualized using a Nikon eclipse 90i microscope with Retiga 2000R (QIMAGING) camera.

### Phosphatidylserine exposure assay

Trypanosomes were reacted with fluorescein isothiocyanate-labeled AnnexinV antibodies (MBL©) and stained with propidium iodide (PI) according to the manufacturer’s instructions. The cells were analyzed by FACS or visualized under a Zeiss LSM 510 META inverted microscope.

### Propidium Iodide (PI) staining for FACS analysis

*T. brucei* cells were fixed in 70% ethanol-30% PBS and stored at 4°C overnight. Cells were then washed once with PBS and incubated on ice for 30 minutes to enable rehydration. The samples were re-suspended in PBS containing 50 μg ml^-1^ RNase A (Roche Diagnostics) for 30 minutes at 4°C and stained with 50 μg ml^-1^ PI (Sigma). Samples and results were analyzed by FACS-Aria and FlowJo (FlowJo, LLC, Ashland, OR), respectively.

## RESULTS

### Silencing of some but not all ER resident proteins induces SLS

SLS was previously shown to be induced in *T. brucei* upon perturbation of functions localized to the ER membrane (8, 9). We subsequently sought to determine whether SLS is induced also by depletion of factors in the ER lumen. To address this question, we silenced key luminal chaperones, namely the *T. brucei* orthologs of *bip* and *crt* mRNAs (43) (44). BiP is an HSP70 chaperone that aids in the translocation and folding of many nascent proteins through cycles of binding and release, whereas CRT is a lectin chaperone of glycoproteins (16).

To perform silencing of the mRNA, cells expressing T7 opposing and stem-loop RNAi constructs were generated. The cells also expressed PTP-tagged proteins integrated at the authentic loci to monitor the degree of silencing (45). All experiments were performed on cloned populations obtained after transfection. Silencing of *bip* and *crt* mRNAs severely inhibited growth (Fig. 1A and B i). The silencing was verified by Northern analysis (Fig. 1A and B ii) and by showing the elimination of the PTP-tagged protein after 2.5 days of silencing (Fig. 1C). Next, the level of SL RNA was examined, and clear and significant reduction of the RNA (∼70%) was observed (Fig. 1C i, ii). SLS is initiated by PK3 phosphorylation leading to its translocation to the nucleus. The phosphorylation of TRF4 then leads to the dissociation of pre-initiation complex and the diffusion of TRF4 in the nucleus (10). To examine the localization of TRF4 and its phosphorylation upon silencing of *bip* and *crt* mRNA, we used antibodies raised against *T. brucei* TRF4. The results clearly demonstrate a shift in the migration of TRF4 upon silencing (Fig. 1D). It was previously demonstrated that this shift results from phosphorylation of TRF4 Ser35 (10). Furthermore, a diffuse localization of TRF4 in the nucleus was also observed in the silenced cells (Fig. 1E i, ii). The data presented were based on imaging of more than 100 cells. The results indicate that as a result of silencing, between 30 to 45% of the cells showed diffuse accumulation of TRF4 in the nucleus, and a significant difference was observed between uninduced and induced cells (Fig. 1E ii). Finally, PCD induced by silencing of *bip* and *crt* mRNAs was examined using the Annexin/PI method. Induction of apoptosis causes externalization of phosphatidyl serine (PS) on the surface of the apoptotic cells. AnnexinV binds to the exposed PS of apoptotic cells. Cells were stained with propidium iodide (PI) distinguish between necrotic and apoptotic cells (46) (see schematic presentation Fig. 1 Fi). Live cells are not stained by either AnnexinV or PI (bottom left panel). During early apoptosis, PS is exposed on the surface, but because the plasma membrane is intact, the cells are not stained with PI (bottom right panel). Late apoptotic or necrotic cells lose their membrane integrity and are stained with both PI and AnnexinV (upper right panel). An early apoptotic population (stained by AnnexinV) was observed after 2 days of silencing, and a late apoptotic population stained with both AnnexinV and PI was observed after 3 days of silencing of both factors (Fig. 1Fii), suggesting that, similar to *sec63* mRNA silencing (7), the silencing of *crt* or *bip* mRNAs induced apoptosis.

**Figure 1.**
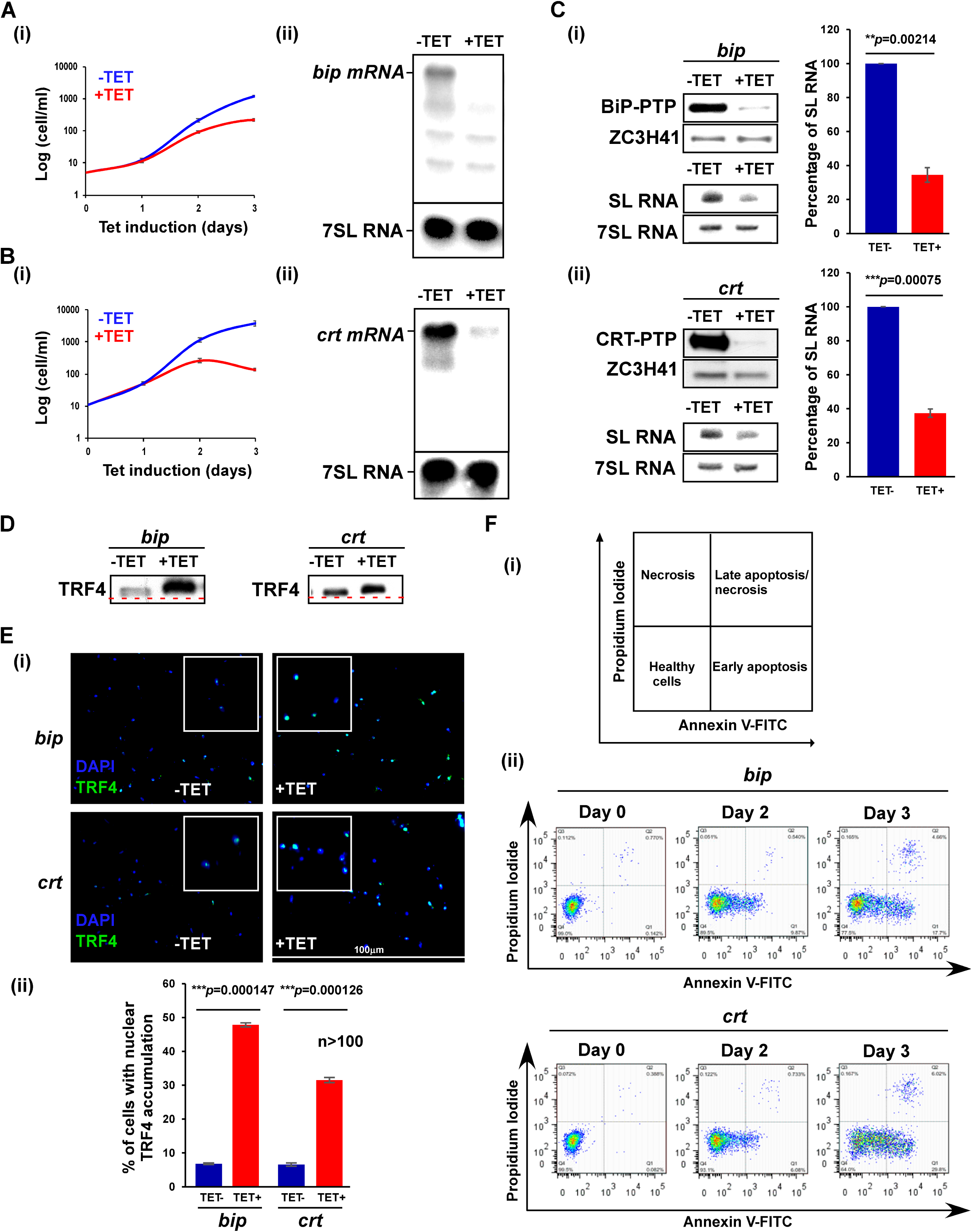
SLS is induced under depletion of factors present in the ER lumen. **A) *bip* mRNA silencing. (i) Growth of cells upon *bip* mRNA silencing.** Uninduced cells carrying the silencing construct (-TET) were compared with cells induced for silencing (+TET) at 27°C for the time period indicated. Data are presented as mean ± S.E.M. Experiments were done in triplicate (n = 3) using the same clonal population. **(ii) Northern analysis.** Total RNA (20 μg) was prepared from cells before (-TET) and after (+TET) *bip* mRNA silencing for 2.5 days, separated on a 1.2% agarose/formaldehyde gel, and subjected to Northern analysis using RNA probes, as indicated. 7SL RNA served as loading control. B) *crt* mRNA silencing was performed in a manner identical to that presented in panel (A). C) The effect of *bip* and *crt* mRNA silencing on SL RNA level. (i) Upper panel. Cells carrying the silencing construct for *bip* and PTP-tagged BiP protein were silenced for the 2.5 days, and the entire cell lysate was subjected to Western analysis with the indicated antibodies. The dilutions used for the antibodies are: IgG (1:10,000) and ZCH341 (1:10,000). **Lower panel**. Cells carrying the silencing construct for *bip* mRNA were silenced for 2.5 days, and the RNA was subjected to Northern analysis with the indicated anti-sense probes. Data are presented as mean ± S.E.M. Experiments were done in triplicate (n = 3) using the same clonal population. *p*-values were determined by Student’s *t*-test **(ii)** The experiment was performed as in (i) but using a cell line expressing the *crt* silencing construct. **D**) *bip* and *crt* mRNA silencing induces the phosphorylation of TRF4. Nuclear extracts were prepared from cells silenced for *bip* or *crt* mRNA for 2.5 days as previously described (27). The proteins were fractionated on 16% SDS-polyacrylamide gel and subjected to Western analysis with anti-TRF4 antibody (diluted 1:10000). The line demonstrates a shift in the migration of the protein in un-induced cells compared to the induced population. **E) Immunofluorescence of cells with anti-TRF4 antibody. (i)** Cells carrying the silencing constructs were subjected to IFA with anti- TRF4 antibody (diluted 1:000), which was detected with anti-rabbit IgG (H+L) conjugated to Alexa Fluor 488 (diluted 1:1000). Cells were visualized by the Nikon eclipse 90i microscope with Retiga 2000R (QIMAGING) camera. Enlargements of the nuclear area are shown as insets. **(ii)** The bar graph represents the quantification of the diffuse pattern of nuclear TRF4 from more than 100 cells per condition. *p*-values were determined using Student’s *t*-test. Data are presented as mean ± S.E.M. **F**) **Analysis of exposed phosphatidylserine on the outer membrane of silenced cells**. **(i)** Schematic representation showing the different cell populations detected by staining with AnnexinV and PI. **(ii)** Cells silenced for the indicated number of days were reacted with FITC-labeled AnnexinV antibody (MBL©) and stained with PI. The identities of the cell lines used are indicated.

After this initial indication that silencing of luminal ER factors mRNAs could induce SLS, we tested whether elimination of additional factors involved in protein folding and quality control in the ER lumen also induce SLS. The factors UGGT, GLU2, and EDEM, which are involved in quality control inside the ER, and ERGIC, which is involved in the transport from the ER to the Golgi, were studied. Cells expressing either T7 RNAi or stem-loop constructs and expressing the PTP-tagged proteins were prepared. Silencing was verified by both Northern and Western blot analyses based on the tagged proteins, and efficient silencing was observed (Fig. 2 A and B). The silencing did not affect cell growth (Fig. 2 C). No difference in the level of SL RNA between uninduced and silenced cells was observed, in contrast to almost 90% reduction in *sec63* silenced cells (Fig. 2Di and ii), suggesting that these factors are not essential for PCF, as opposed to bloodstream form trypanosomes (26).

**Figure 2:**
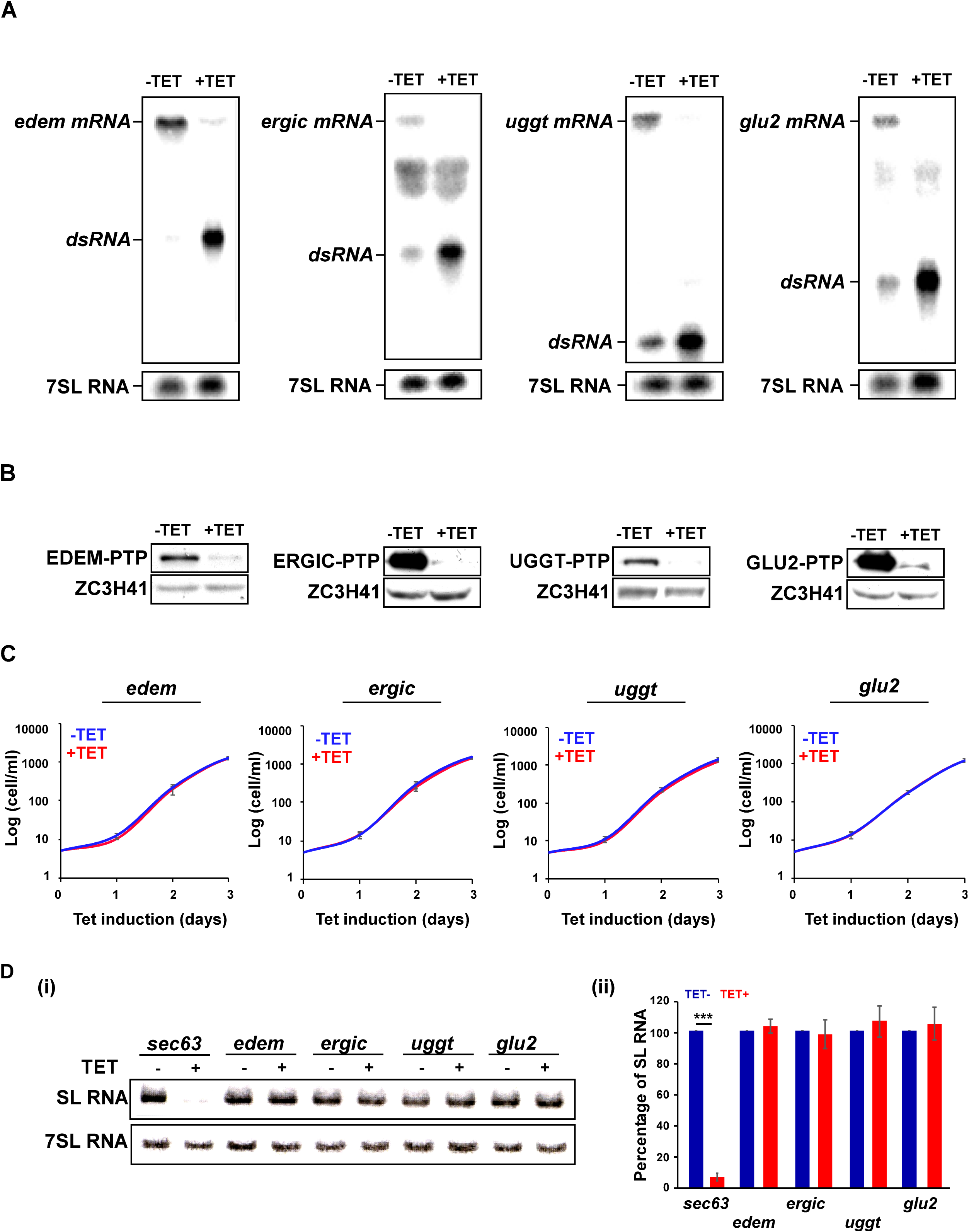
Silencing of *uggt, glu2, edem,* and *ergic* mRNAs did not affect PCF growth or induce SLS. **A) Northern analysis.** Total RNA (20 μg) was prepared from cells before (-TET) and after 2.5 days silencing (+TET), separated on a 1.2% agarose/formaldehyde gel, and subjected to Northern analysis using RNA probes, as indicated. 7SL RNA served as loading control. The silenced mRNA and dsRNA positions are indicated. **B) Western analysis demonstrating the silencing of the mRNA**. Western analysis was performed after 2.5 days of silencing demonstrating the depletion of the tagged protein. The dilutions used for the antibodies are: IgG (1:10,000) and ZCH341 (1:10,000). **C) Silencing of the genes has no effect on cell growth.** Uninduced cells carrying the silencing construct (-TET) were compared with cells induced for silencing (+TET) at 27°C during the time period indicated. Data are presented as mean ± S.E.M. Experiments were done in triplicate (n = 3) using the same clonal population. **D) SL RNA level upon silencing. (i)** Cells carrying the silencing construct for the indicated mRNAs were silenced for 2.5 days, and the RNA was subjected to Northern analysis with the indicated anti-sense probes. Data are presented as mean ± S.E.M. Experiments were done in triplicate (n = 3) using the same clonal population. *p*-values were determined by Student’s *t*-test. **(ii)** The bar graph represents the quantification of SL RNA upon silencing of the indicated mRNAs. Data are presented as mean ± S.E.M. Experiments were done in triplicate (n = 3) using the same clonal population. *p*-values were determined by Student’s *t*-test

### Depletion of the sulfhydryl oxidases ERO1 and QSOX1 induces SLS

We next asked whether interfering with secretory pathway functions other than chaperoning and quality control could induce SLS. Another feature of the secretory pathway is its ability to promote formation of disulfide bonds during protein folding and assembly. Mammalian ERO1 catalyzes disulfide bond formation in the ER. Another catalyst of disulfide formation, QSOX1, is localized to the Golgi apparatus or secreted from mammalian cells (47). Antibodies raised against *T. brucei* QSOX1 were used to establish the localization of this enzyme in the parasite. Co-localization of QSOX1 was examined with the Golgi network (TGN) protein marker GRIP70 (48), the Golgi stack marker GRASP (49), and SEC24, which marks the budding of COPII vesicles from the ER (48). The results (Fig. 3A) show closest co-localization of *T. brucei* QSOX1 with GRASP, indicating that QSOX1 is Golgi-localized in *T. brucei* as in mammalian cells.

**Figure 3:**
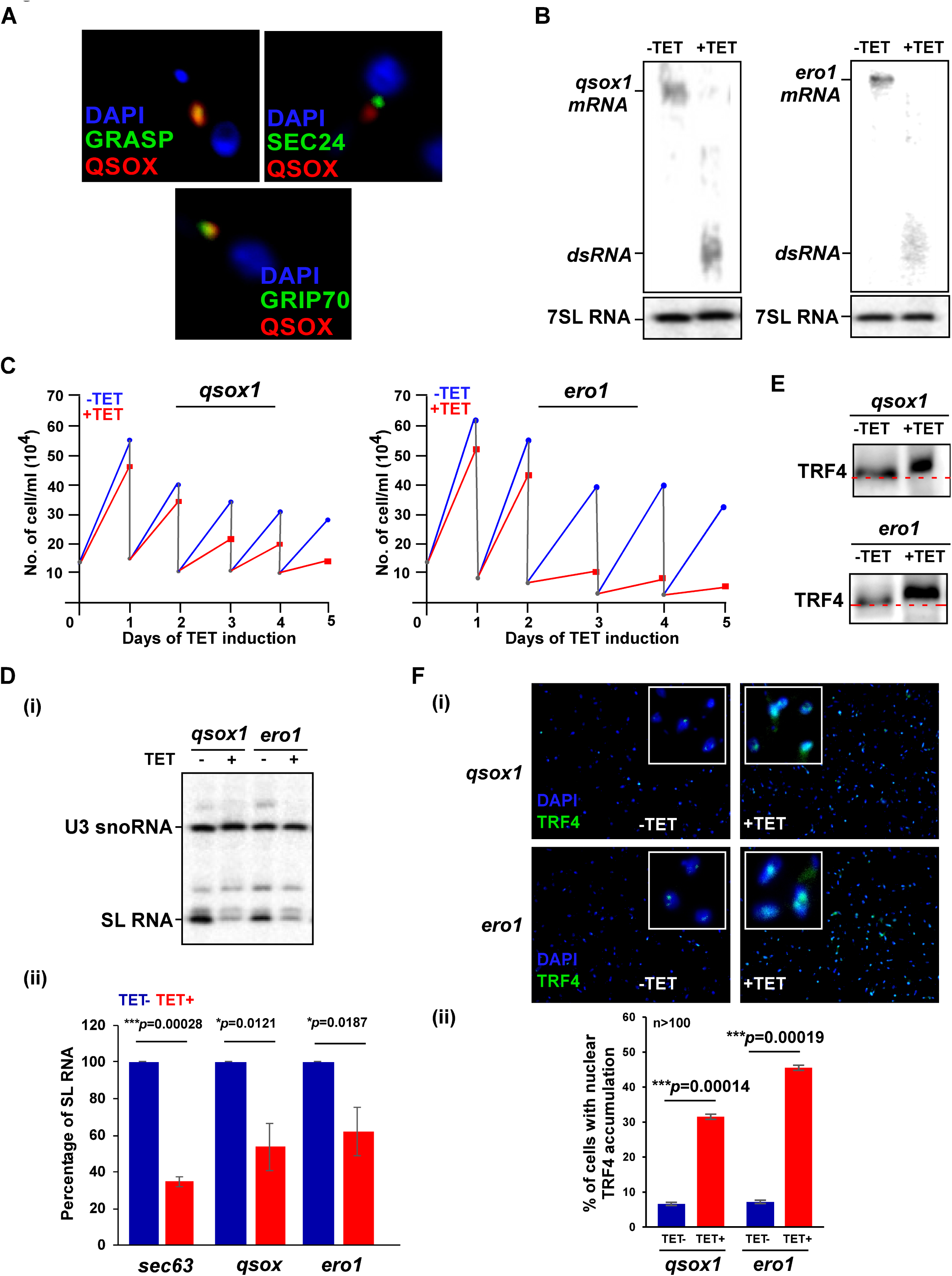
The silencing of *qsox1* and *ero1* mRNA induces SLS. **A) Localization of QSOX1.** PCF cells were subjected to IFA with rat anti-QSOX1 antibody, which was detected using anti- rat IgG conjugated to Alexa Fluor 488 (diluted 1: 10000) as well as with either rabbit GRASP, GRIP70 or SEC24 antibody (48, 49) (diluted 1:1000), and detected with anti-rabbit IgG conjugated to Cyc3. Cells were visualized by the Nikon eclipse 90i microscope Retiga 2000R (QIMAGING) camera. Nuclei were stained with DAPI. **B) Northern analysis.** Cells expressing the stem-loop silencing constructs for *qsox1* and *ero1* mRNA were induced for 2.5 days, and RNA was subjected to Northern analysis with their gene-specific RNA probes. The positions of *qsox1*and *ero1* mRNA and dsRNA are indicated. **C) *qsox1* and *ero1* silencing induces growth arrest.** The growth of cells prior to and after tetracycline addition was compared. Both un-induced and induced cultures were diluted daily to 2 x 10^4^ cells per ml. **D). The effect of *qsox1* and *ero1* mRNA silencing on SL RNA**. **(i)** Total RNA (10µg) was prepared from *qsox1* and *ero1* mRNA silenced cells after 2.5 days of induction, and was subjected to primer extension with anti-sense SL RNA oligo. The level of U3 was used to control for equal loading. The samples were fractionated on 6% polyacrylamide/7M urea gels. **(ii)** The bar graph represents the quantification of SL RNA upon silencing of the indicated mRNAs. Data are presented as mean ± S.E.M. Experiments were done in triplicate (n = 3) using the same clonal population. *p*-values were determined by Student’s *t*-test. **E) *qsox1* and *ero1* mRNA silencing induces the phosphorylation of TRF4.** Nuclear extracts were prepared from cells after 2.5 days of silencing, as previously described (27). The proteins were fractionated on 16% SDS-polyacrylamide gel and subjected to Western analysis with anti-TRF4 antibody (diluted 1: 10000). The line demonstrates a shift of protein migration in un-induced cells compared to the induced population. **F) Immunofluorescence of QSOX1 and ERO1 cells with TRF4 antibody. (i)** Cells carrying the silencing constructs either un-induced or after 2.5 days of silencing were subjected to IFA with TRF4 antibody (diluted 1:000), which was detected with anti-rabbit IgG (H+L) conjugated to Alexa Fluor 488 (diluted 1:1000). The nucleus was stained with DAPI. Cells were visualized under the Nikon eclipse 90i microscope with Retiga 2000R (QIMAGING) camera. Enlargements of the nuclear area are shown as insets. **(ii)** The bar graph represents the quantification of diffuse nuclear TRF4 from more than 100 cells per condition. *p*-values were determined by Student’s *t*-test. Data are presented as mean ± S.E.M.

Silencing of either *qsox1* or the *ero1* sulfhydryl oxidase mRNAs (Fig. 3B) indicates that these factors are essential for growth (Fig. 3C). The level of SL RNA was examined by primer extension, and reduction of about 50% of SL RNA was observed (Fig. 3D i, ii). Primer extension analysis was used to demonstrate that SL RNA is not degraded under SLS, suggesting that the reduction in SL RNA level stems from SL RNA transcription shut-off. The phosphorylation of TRF4 (Fig. 3E) demonstrates retardation of the TRF4 upon silencing. Next, the distribution of TRF4 in the nucleus was examined upon silencing. Silencing resulted in diffuse accumulation of TRF4 in the nucleus compared to uninduced cells (Fig. 3F i and ii). These results show that that depletion not only of factors localized on the ER, but also in the ER lumen as well as outside the ER can induce SLS.

### Depletion of TIMRHOM1 affects protein translocation across the ER, mitochondrial function, and induces SLS

The proteome analysis of cells in which SLS had been induced by silencing of *sec63* indicated an increase in the level of TIMRHOM1 protein (27). To verify this elevation, the level of the protein was examined upon SLS induced by *sec63* silencing, and a four-fold increase was observed (Fig. 4A). Thus, TIMRHOM1 appears to be connected to the SLS pathway. Because of the increase in TIMRHOM1 during *sec63* silencing, we suggested that its induction may assist in coping with protein translocation defects across the ER. To test the effect of depleting TIMRHOM1, a stem-loop construct was used for silencing, which was verified by Northern analysis (Fig. 4B). We also co-silenced *sec63* with *timrhom1* to examine if *timrhom1* mRNA silencing exacerbates the protein translocation defects across the ER. The co-silencing was confirmed by Northern analysis (Fig. 4C). To examine the effect of the co-silencing on protein translocation across the ER, the levels of two proteins, VH+-ppase, which is a membrane protein located in the acidocalcisome, and the glycosylphosphatidylinositol (GPI)-anchored protein EP, both affected by *sec63* silencing (8), were examined. To our surprise, since TIMRHOM1 is localized to the mitochondria (28), a clear reduction in the levels of both ER translocated proteins was observed upon *timrhom1 mRNA* silencing (Fig. 4D). Co-silencing did not worsen the defects (Fig. 4D). To verify that, indeed, TIMRHOM1 affects mitochondrial integrity and function, mitochondria were stained with mitotracker, and mitochondrial function was assessed using Tetramethyl rhodamine methyl-ester (TMRM), which accumulates in the negatively charged active mitochondria, and reports on mitochondrial membrane potential (ΔΨm). Mitotracker staining showed a single long mitochondrion in uninduced cells, which upon silencing, appeared fragmented (indicated by arrows) (Fig. 4Ei). A significant difference in fragmentation was detected upon silencing (Fig. 4Eii). The reduction in ΔΨm was more rapid in *timrhom1* mRNA silenced cells compared to *sec63* mRNA silenced cells (Fig. 4F), as expected because TIMRHOM1 is a mitochondrial protein involved in protein translocation and the effect on mitochondria is primary, whereas the effect of *sec63* silencing on the mitochondria is likely to be secondary.

**Figure 4:**
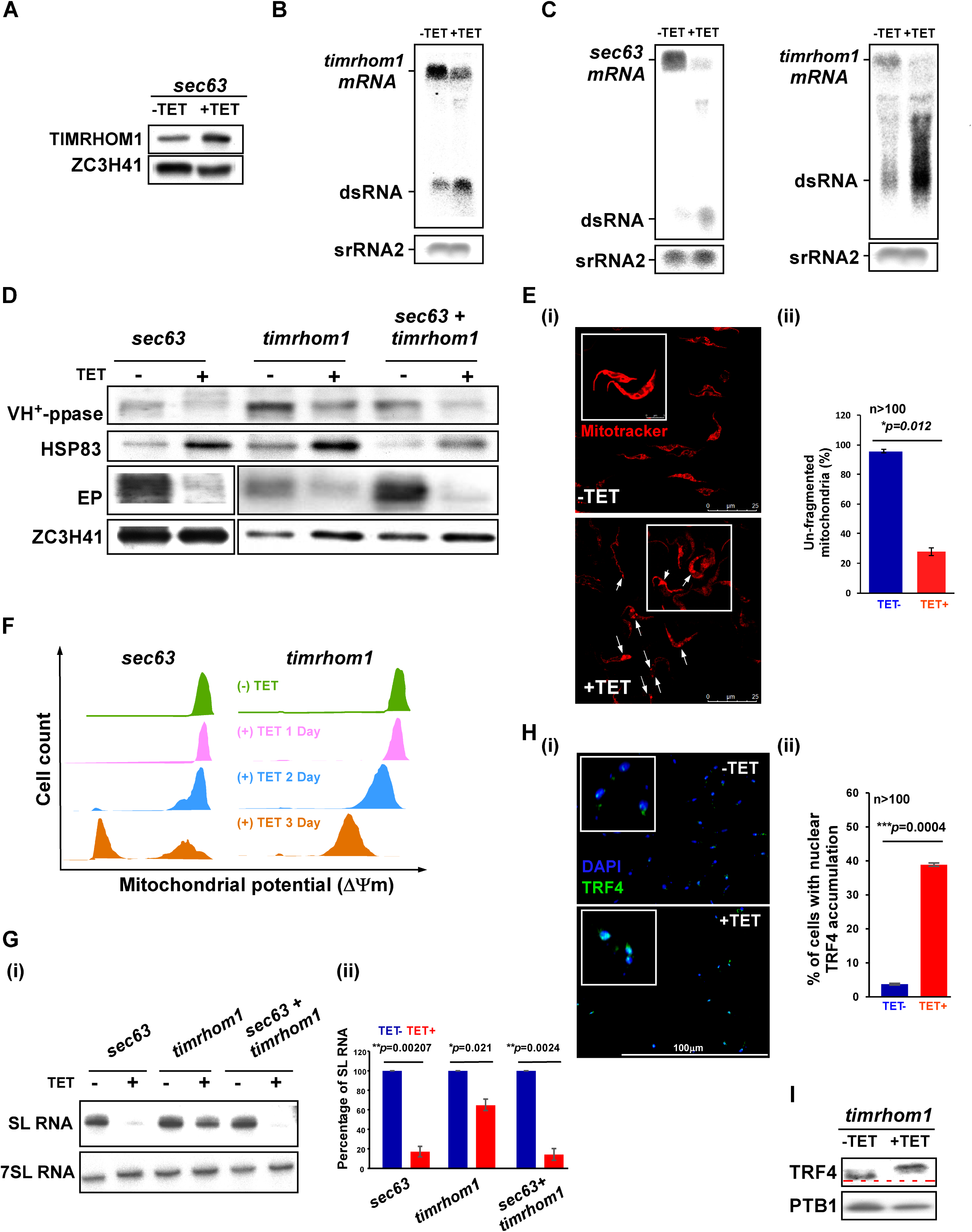
Silencing of *timrhom1* mRNA induces SLS. **A) The level of TIMRHOM1 upon SLS induction.** Cells carrying the silencing construct for *sec63 mRNA* were silenced for 2.5 days, and the whole cell lysate was subjected to Western analysis with the indicated antibodies. The dilutions used for the antibodies were: a) TIMRHOM1 (1:500) and ZCH341 (1:10,000). **B) Silencing of *timrhom1* mRNA**. Cells expressing the stem-loop construct for silencing *timrhom1 mRNA* were induced for 2.5 days, and RNA was subjected to Northern analysis with the gene- specific RNA probes, and srRNA2 was used to control for equal loading. The positions of the mRNA and dsRNA bands are indicated. **C) Co-silencing of *sec63* and *timrhom1* mRNA**. Northern blot analysis was performed on RNA from cells carrying the silencing constructs for both *sec63* and *timrhom1 mRNA.* The blots were probed separately with gene-specific probes, and srRNA2 was used to control for equal loading. **D) Effect of *timrhom1 mRNA* silencing on protein translocation across the ER**. Proteins from *sec63*, *timrhom1*, and *sec63/ timrhom1 mRNA* cells silenced for 2.5 days were subjected to Western analysis and reacted with antibodies to anti-vacuolar VH+ppase antiserum (diluted 1:5000) as well as anti-EP mAb 247 (diluted 1:10,000). **E**) **Silencing of *timrhom1 mRNA* induces mitochondrial fragmentation. (i)** Cells expressing the silencing construct for *timrhom1 mRNA* were silenced for 2.5 days and stained with mitotracker. Enlargements of cells are shown in the insets. Cells with fragmented mitochondria are marked with white arrows. **(ii)** The bar graph represents the quantification of unfragmented mitochondria from more than 100 cells per condition. *p*-values were determined by Student’s *t*-test. Data are presented as mean ± S.E.M. **F) Silencing of *timrhom1* mRNA induces loss of mitochondrial membrane potential.** Cells were harvested and loaded with 150 nM TMRM in serum free medium. The samples were incubated in the dark for 15 minutes at 27 °C and then analyzed by FACS. The cell count of cells along with ΔΨm is plotted comparing the cells before and during the first 3 days of silencing. The data compare *sec63* silenced cells to that of *timrhom1* mRNA silencing. **G) *timrhom1* mRNA silencing reduces the level of SL RNA**. **(i)** Total RNA (10µg) was prepared from *sec63*, *timrhom1,* and *sec63* and *timrhom1* silenced cells after 2.5 days of induction, fractionated on 10% polyacrylamide/7M urea gel, and subjected to Northern analysis with SL RNA and 7SL RNA probes. **(ii)** The bar graph represents the quantification of SL RNA upon silencing of the indicated mRNAs. Data are presented as mean ± S.E.M. Experiments were done in triplicate (n = 3) using the same clonal population. *p*-values were determined using Student’s *t*-test. **H) Changes in TRF4 localization upon *timrhom1 mRNA* silencing. (i)** Cells carrying the *timrhom1 mRNA* silencing construct were induced for 2.5 days and subjected to IFA with TRF4 antibody (diluted 1:000), which was detected with anti-rabbit IgG (H+L) conjugated to Alexa Fluor 488 (diluted 1:1000). The nucleus was stained with DAPI. Cells were visualized by the Nikon eclipse 90i microscope with Retiga 2000R (QIMAGING) camera. Enlargements of the nuclear area are shown as insets. **(ii)** The bar graph represents the quantification of diffused pattern of nuclear TRF4 upon silencing from more than 100 cells per condition. *p*-values were determined by Student’s *t*-test. Data are presented as mean ± S.E.M. **I) *timrhom1* mRNA silencing induces the phosphorylation of TRF4.** Nuclear extracts were prepared from cells after 2.5 days of silencing, as previously described (27). The proteins were fractionated on 16% SDS-polyacrylamide gel and subjected to Western analysis with anti-TRF4 antibody (diluted 1: 10000). The line demonstrates the shift of protein migration in un- induced cells compared to the induced.

Because *timrhom1* mRNA silencing induced mitochondrial fragmentation, as in SLS induction, we examined whether *timrhom1* mRNA silencing itself induces SLS. Indeed, Northern analysis based on three replicates showed that the level of SL RNA was reduced upon *timrhom1* mRNA silencing but to a lesser extent compared to *sec63* silencing (Fig. 4Gi and ii). Immunofluorescence of TRF4 in the nucleus demonstrates the massive accumulation of TRF4 upon *timrhom1* mRNA silencing (Fig. 4Hi and ii). In addition, Western analysis with anti-TRF4 (Fig. 4I) showed that silencing of *timrhom1* mRNA induces the phosphorylation of TRF4. Thus, SLS induction is not restricted to perturbation of only secretory pathway compartments but can also be induced by defects in the mitochondria due to blockage of protein translocation.

### PK3 undergoes phosphorylation under SLS, which affects its translocation to the nucleus

The serine/threonine family eIF2 kinases such as PERK undergo auto-phosphorylation upon activation (50–52). PK3 belongs to this family of kinases (53) and hence is expected to undergo auto-phosphorylation upon activation. The activation of PERK is followed by the autophosphorylation of the kinase domain, which provides PERK with full catalytic activity (54). In the crystal structure of the PERK kinase domain, the N-terminal lobes are responsible for dimerization, forming homodimers constituted by two back-to-back monomers. Following autophosphorylation at the activation loop (C-lobe), the structure becomes ordered and is partially stabilized and able to bind its substrate (55).

We have so far shown that different cues induce SLS including low pH, chemicals such as dithiothreitol (DTT) and 2-deoxy-D-glucose (2-DG), and depletion of factors involved in protein translocation into the ER (7–9), as well as factors involved in protein folding or assembly localized in the ER lumen, or even the Golgi apparatus or mitochondria, as demonstrated here. It is currently unknown whether all these cues activate PK3 in the same manner. To answer this question, we first identified the protein modifications that PK3 undergoes during SLS induced by *sec63* silencing. To this end, we used the cell line expressing both the *sec63* silencing and tagged PK3-PTP constructs. The purification to extract the majority of the kinase required several detergents, as specified in Materials and Methods, and only under these conditions was it possible to release PK3 from its association with membranes. The PTP-tagged protein was purified as previously described (10) and selected on a titanium oxide column to enrich for phosphorylated proteins. Alternatively, the purified PK3 was separated on a 10% SDS-polyacrylamide, purified from the gel, and subjected to mass-spectrometry. The detection of PK3 after the detergent extractions and the quality of purification of the PK3 before and after *sec63* silencing is presented in Fig. 5A and B, respectively. The list of the phosphorylation sites revealed by these two purification methods is presented in (Table S2). The phosphorylation sites induced by SLS on the PK3 sequence are presented in Fig. 5C, which shows that four of the phosphorylation sites are clustered in the kinase domain.

**Figure 5:**
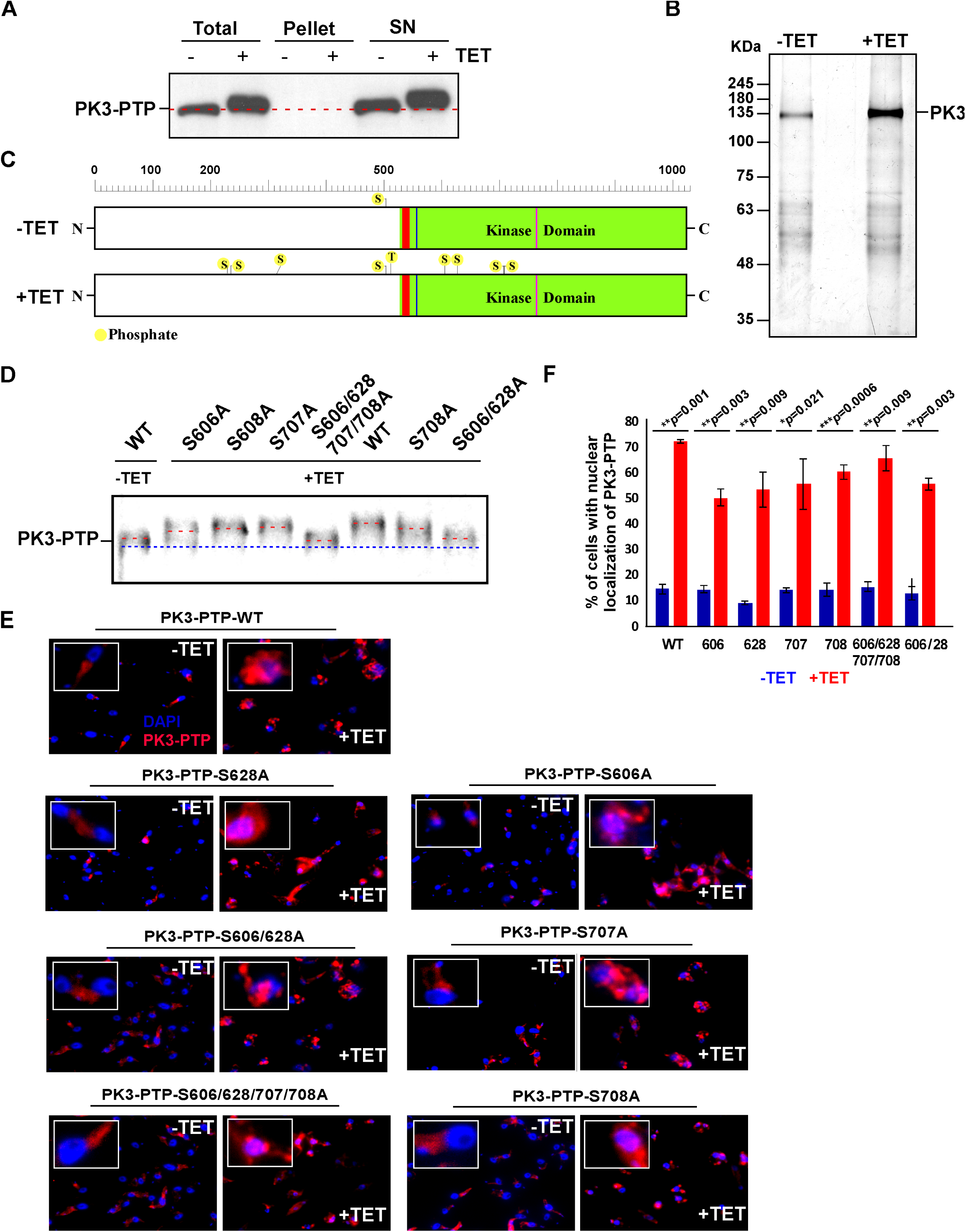
PK3 phosphorylation under SLS. **A) Purification of PK3-PTP protein from SLS-induced cells.** Extract was prepared from 10^10^ silenced cells after treatment with detergent rich buffer as detailed in Materials and Methods. Aliquots from total cell extracts, soluble (SN) and insoluble material (1/50) were subjected to Western analysis. **B) Purified PK3.** The purification was performed as previously described (33) from the extract in A, above. The eluted material (1/10) was fractionated on a 10% SDS-polyacrylamide gel and stained with silver. The remaining material was fractionated on a gel, eluted and subjected to MS (Table S2) **C) Schematic presentation of the localization of the sites that undergo phosphorylation under SLS**. The kinase domain (positions 528-1022) is depicted in green, and the ATP binding domain (positions 534-542) is in red. **D) Mutations in the phosphorylation sites and their effect on PK3 modification.** The mutations were constructed as described in Materials and Methods. Cells carrying the *sec63* mRNA silencing construct and the different PK3 mutations were subjected to Western analysis with Rabbit-IgG serum (Sigma) followed by reaction with anti-rabbit IgG (H+L) conjugated to horseradish peroxidase. The positions of the mutations are indicated, and the line shows the difference in migration of the wild-type tagged protein and the PK3 mutants. **E) Immunofluorescence of PK3 upon induction of SLS.** Cells expressing the *sec63* silencing construct and PK3-PTP mutations were silenced for 2.5 days. The cells were subjected to IFA by first reacting the cells with Rabbit- IgG serum (Sigma) (diluted 1:1000), and then with anti-rabbit IgG (H+L) conjugated to Alexa Fluor 488 (diluted 1:1000). The nucleus was stained with DAPI. Enlargements of the nuclear area are shown as insets. **F)** The bar graph represents the quantification of nuclear localization of PK3-PTP upon *sec63* silencing from more than 100 cells per condition. *p*-values were determined by Student’s *t*-test. Data are presented as mean ± S.E.M. The identity of each mutation is indicated.

To verify the MS results and demonstrate that these sites are indeed phosphorylated as a result of SLS induction, single, double as well as quadruple mutations were introduced at the following positions in PK3: 606, 628, 707, 708, 606/628 and 606/628/707/708. The serine to alanine mutant constructs were introduced into cells carrying the *sec63* silencing construct. The results demonstrate that the mutated positions are indeed phosphorylated under SLS, since upon induction of SLS, the mutated tagged protein migrated differently than the wild-type tagged PK3 (Fig. 5D). Retardation of migration is consistent with the number of mutated residues. Only the mutant in which four resides were mutated exhibited a defect in its ability to undergo phosphorylation. Next, we examined whether the mutated proteins could translocate to the nucleus. To this end, we examined the localization of the mutated proteins by immunofluorescence using an antibody that recognizes protein A present in the PTP tag. The results (Fig. 5E and F) demonstrate that translocation to the nucleus of the different mutations was compromised and was less efficient when compared to the wild-type PK3. However, since PERK-like kinases function as dimers (55), it is possible that the mutants formed heterodimers composed of wild-type and mutant PK3 monomers, which could still translocate to the nucleus.

To examine if complete abrogation of phosphorylation would completely eliminate the nuclear localization under SLS, we performed structural modeling of the PK3 kinase domain (KD), to identify residues that could potentially inactivate the kinase function. The homology model of PK3 (UniProtKB identifier Q583N6) was generated based on the crystal structures of the homologous proteins: human PERK (PDB ID: 4G31) (56) and mouse PERK (PDB ID: 3QD2) (55) KD domains. Sequence alignment of the entire KD domains showed 14.6% sequence identity between PK3 and its homolog (Fig. S1). However, since the template structures mentioned above lack many residues present in PK3, removing the corresponding missing residues from the query sequence of PK3 increased the sequence identity to 26.9%. The homology model for PK3 KD was produced by the MODELLERv.9.2 program (57, 58), and further side chain refinement was performed using Predict Side Chains tool as implemented in the Maestro program (Schrödinger, USA). The resulting homology model (Fig. 6A) was found to have favorable stereochemical qualities, with 96.3% of residues residing in the most favored regions and additionally allowed regions of the Ramachandran plot, and an overall G-factor of -0.14. The model’s Prosa profile was similar to the template, with a z-score of -4.76 (59). Template/target root mean square deviation of atomic positions (RMSD), was calculated to determine the similarity between the two protein structures. The smaller the RMSD, the more similar the structures. Values based on Cα of the backbone atoms were 2.1Å and 2.4Å for PK3 model / human PERK (56) and PK3 model / mouse PERK (55), respectively (Fig. 6A). All these parameters support the quality of the proposed model. The model of PK3 KD (red) is shown in Fig. 6A and demonstrates high similarity to the two templates (human PERK (depicted in green) and mouse PERK (depicted in blue)). The functional domains are indicated. After optimizing the model of PK3 KD, the ATP binding domain was predicted and was found to have 63.6% identity with the homologs (Fig. S1). The phenylalanine Phe771 (F771) (indicated in Fig. 6B) present in the ATP binding domain was found to be conserved among the sequences used for building the homology model (Fig.6B) (i.e., Phe943 in 4G31 of human, and Phe942 in 3QD2 of mouse PERK). This Phe771 is likely to contribute to the binding of ATP. In addition, the crystal structure of the complex between the PERK inhibitor GSK2606414 and the human PERK protein reveals that the ligand forms a π interaction with Phe943 in the protein’s ATP binding site, supporting the role of this residue in ATP binding (56). Finally, docking using the Glide SP protocol (60, 61) of ATP into the ATP binding site of the PK3 model resulted in two T-shaped π−π interactions between the adenosine moiety of ATP and the conserved Phe771 (Fig. 6B marked with dashed arrows). Taken together, these observations suggested that mutating Phe771 may prevent PK3 from binding ATP thereby rendering the protein non-functional. To examine if this is indeed the case, a mutation was introduced in this position converting phenylalanine to leucine in the C-terminal MYC-tagged version of the gene. The gene was integrated *in-situ* in PCF cells and induction of SLS was performed by low pH 5.5 (9). Upon induction of SLS at low pH, the wild-type MYC-PK3 migration was retarded, suggesting that the MYC tagged protein was functional (Fig. 6C lanes 1, 2). However, the MYC tagged mutated protein was unable to undergo phosphorylation (Fig. 6C lanes 3, 4). Next, we examined the translocation of the protein following SLS induction by low pH (Fig. 6Di and ii). The results show dramatic effect of this mutation on the ability of the mutated protein to translocate to the nucleus, (only 30%) (Fig. 6Dii). The residual translocation could be explained by formation of dimers with a wild-type monomer. However, the marked reduction in the translocation to the nucleus of the non-phosphorylated protein upon conditions that induce SLS suggest that PK3 phosphorylation is essential for its translocation.

**Figure 6.**
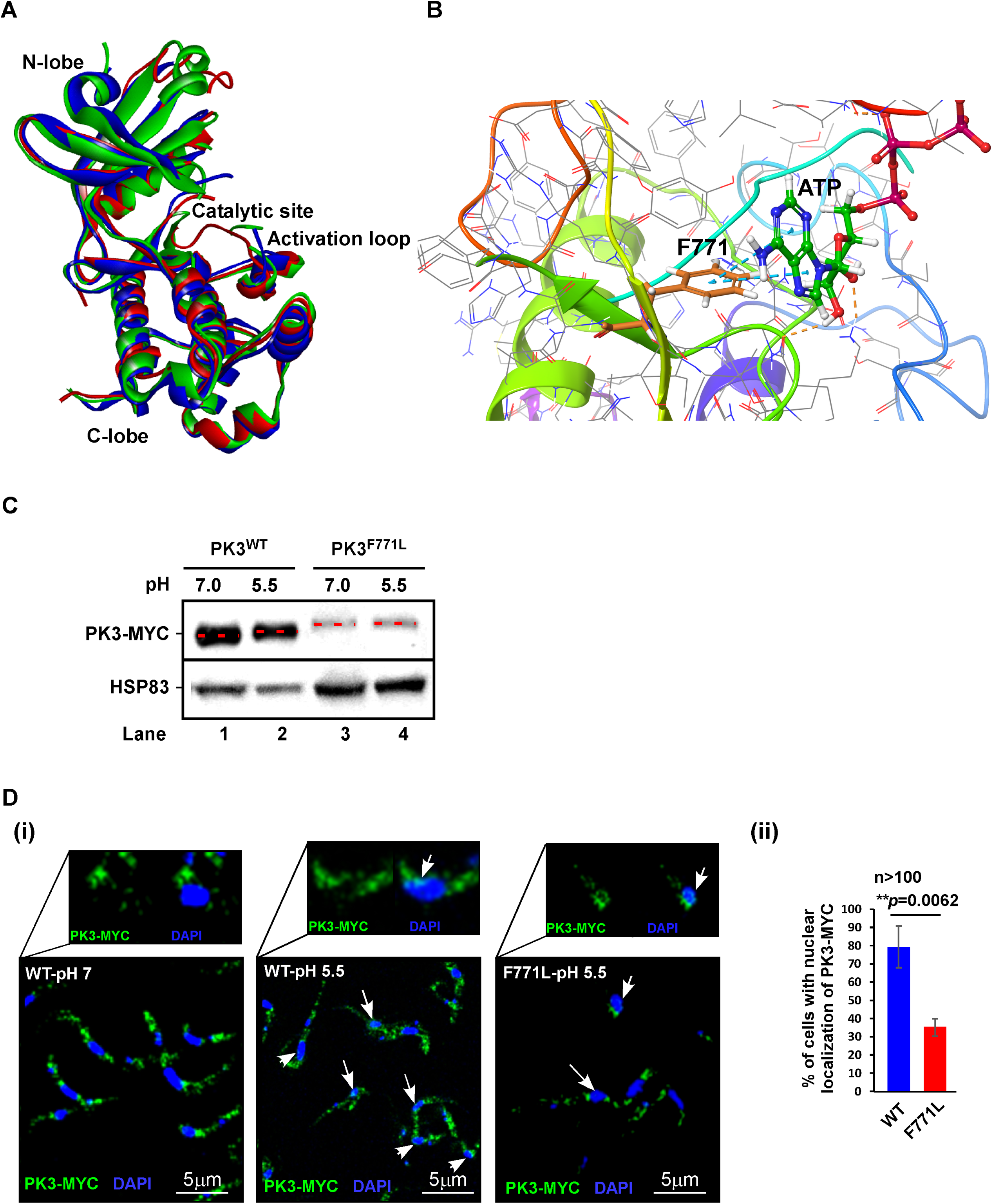
Modeling PK3 KD domain. **A) Homology model of PK3 KD superimposed on the crystal structures of human and mouse PERK KD.** The modeling was based on the human PERK (56) and mouse PERK (55). The structures are shown as ribbon diagrams, and colored in red, green, and blue, representing PK3, human and mouse proteins, respectively. The position of the N-lobe, C-lobe, catalytic site, and activation loop are indicated. **B) Docking of ATP in its binding site on the proposed PK3 model.** The Phe 771 (F771) that makes contact with the adenosine moiety of ATP via two T-shape π−π interactions is indicated. The interactions are illustrated with blue dashed lines. **C) PK3^WT^-MYC tagged protein but not PK3^F771L^ undergoes phosphorylation under SLS.** Cells expressing the PK3 tagged with MYC were incubated at either pH 7.0 (control) or at pH 5.5 for 3 days to induce SLS as previously described (9). Whole cell lysates were prepared and subjected to Western analysis with anti-MYC antibody (9E10, Santa-Cruz). HSP83 was used to monitor protein levels. The lanes are numbered. **D) Changes in the localization of PK3-MYC upon SLS induction. (i)** Cells were incubated at the indicated pH as described above and subjected to IFA with anti-MYC antibodies. The nuclei were stained with DAPI. Enlargement of a portion of the images is shown in the inset. The cells harboring the tagged PK3-MYC in their nucleus are indicated with arrows. **(ii)** Bar graph representing the quantification of nuclear localization of PK3-MYC at pH 5.5 from more than 100 cells per condition. *p*-values were determined by Student’s *t*-test. Data are presented as mean ± S.E.M. The identity of each mutation is indicated.

## DISCUSSION

In this study, we demonstrate that SLS is induced by depletion of protein factors including not only those located in the ER membrane, but also factors inside the ER and involved in protein folding (BIP, CRT) and protein oxidative modification (EROI). SLS was also found to be induced by depletion of an enzyme catalyzing disulfide bond formation located in the Golgi apparatus (QSOX1). The most surprising finding was that SLS is also induced upon depletion of TIMRHOM1, a factor involved in protein translocation into the mitochondria (28), supporting the role of PK3 as the kinase that regulates homeostasis between the ER and the mitochondria.

Two *T. brucei* iRhomboid proteins exist, TIMRHOM1 and TIMRHOM2, and were shown to be genuine components of the inner mitochondrial import system and to associate with a protein translocation intermediate stalled on its way into the mitochondria (28). However, the level of only TIMRHOM1 was elevated in SLS-induced cells (27). In mammals iRhomboid proteins, especially from the DERLIN family, were shown to be associated with the ER and directly involved in ERAD (62). The fact that iRhomboid proteins lack proteolytic activity but can bind and stabilize unfolded domains makes them most suitable to regulate the fate of membrane proteins (63). However, it was recently suggested that the two trypanosome Rhomboid-like proteins are functional homologues of the yeast pre-sequence translocation pore to the mitochondria. Of note, the trypanosome Rhomboid-like proteins are not related to the eukaryotic iRhomboid proteins (DERLIN, PARL) nor to their bacterial counterparts, and thus may have originated from the endosymbiont that gave rise to the mitochondrion (28).

The observation that PK3 is phosphorylated and trafficked upon depletion of ER factors and the mitochondrial TIMRHOM1 suggests that it may respond to perturbations taking place either in the ER or mitochondria. If so, it is possible that this kinase is localized to ER-mitochondria contact sites. An ER-mitochondria encounter structure (ERMES) was described and shown in yeast to be important for lipid transport and calcium signaling (64). In mammalian cells, the mitochondria-ER contact sites (MERC) mediate the interaction between the mitochondrial porin-voltage-dependent anion selective channel protein 1 (VDAC1) and the ER Ca^2+^ channel (inositol 1,4,5- trisphosphate receptor IP(3) R) (30). It is unclear at the present time what type of ERMES exists in trypanosomes. A functional homologue of Mdm10 that is part of the yeast ERMES was identified in *T. brucei* and named TAC40. TAC40 mediates the linkage between mitochondrial DNA and the basal body connecting the mitochondrion with the flagellum of the parasite. However, no similar ERMES mediating the contact between the ER and the mitochondria and related to the yeast complex has been identified in trypanosomes (65). In addition, the Ca^2+^ channel IP(3) R, which is part of mammalian ER contact sites (30), localizes to trypanosome acidocalcisomes, which is the organelle mediating Ca^2+^ storage and bearing membrane contact sites with the mitochondria (66). Thus, we still do not know which trypanosome proteins mediate the contact between the ER and mitochondria. We are currently identifying the proteins associated with PK3 under normal conditions and upon SLS induction, hoping to identify proteins that contact PK3 and constitute the ER-mitochondria contact sites. In mammalian cells, the PERK kinase controls mitochondria function by attenuating mitochondrial protein import in response to stress, thereby preventing the accumulation of damaged or non-native proteins that can interfere with mitochondrial function in response to ER stress (67). In addition, PERK induces up-regulation of the level of stress-induced mitochondrial hyperfusion (SIMH), a protective mechanism that suppresses pathologic mitochondrial fragmentation and promotes mitochondrial functions such as ATP production, which helps prevent mitochondrial fragmentation (68).

The finding that depletion of proteins involved in ER quality control does not induce SLS in PCF is intriguing, and indicates that these factors may not be necessary for the survival of PCF trypanosomes. Indeed, stress-induced misfolding occurs along the life cycle of these parasites. The non-dividing transmissive forms of the parasite, short stumpies (mammals to insect) and metacyclics (insect to mammals) may require ERAD capacity for cellular re-modelling upon transmission. BSF ERAD is presumably critical because of antigenic variation. During antigenic variation, different versions of VSG can be formed but many versions are likely to be misfolded. There must therefore be a system with “disposal capacity” to cope with the stress of misfolded VSG to be able to survive catastrophic switches. ERAD may provide the mechanism to cope with such unsuccessful switches (69). Indeed, ERAD was demonstrated in trypanosomes when a mutated version of transferrin receptor accumulated in the ER and was degraded by the proteasome (69). Thus, ERAD functions, and possibly all the factors that were studied here that did not induce SLS such as UGGT, GLU2 and EDEM, are not essential in PCF grown under normal conditions. However, these factors were shown to be essential in BSF, supporting the crucial role of ERAD at this life stage (26) as stated above.

A recent study suggested that SLS induction is a stage-specific pathway, as *sec63* silencing did not induce SLS in BSF (70). However, the silencing in that study was done for a relatively short time (24 hrs), which is likely insufficient for generating defects in protein translocation necessary to induce SLS. The studies in PCF were always performed under longer silencing periods (2-3 days) (7, 9). We have previously shown that *sec63* silencing in BSF induces SLS after at least 2.5 days (7).

SLS is induced following treatment with compounds known to induce ER stress in other eukaryotes (7) or exposure to low pH (9). However, SLS is only induced following depletion of some essential factors (71). For example, SLS is not induced when depleting the SRP protein SRP54, despite being essential for viability (72). Depletion of SEC71, affecting the biogenesis of signal peptide (SP)-containing proteins but not polytopic membrane, also did not induce SLS (73). Note that silencing of factors involved in *trans*-splicing results in the accumulation of SL RNA and its secretion from the cell (71). Heat-shock in both life stages of the parasite also induces an increase in the level of SL RNA and its secretion. SLS induction is a point of no return and represents a decision taken by the parasite to initiate a death pathway.

This study demonstrates that PK3, like other eIF-2α kinases, undergoes phosphorylation on multiple sites. To this end we used PhosphoSitePlus database (http://www.phosphosite.org) (74) and compared the phosphorylation sites of human and mouse PERK with those observed on PK3 (Fig S2). To be able to evaluate the functional importance of the modified positions, we compared the kinase domain of PERK to the trypanosome homologues of eIF-2α (PK1-2) as previously described (53). We also compared all the reported phosphorylation sites on the different PERK kinases (75–92) to that of PK3 (Fig S2). We noticed that Y554 of PK3 is analogous to the phosphorylated Y619 in the human kinase. Despite identifying 23 peptides covering this residue, there was no evidence of its phosphorylation under SLS activation. Failure to detect such a phosphorylation site does not imply that it does not exist. Phosphorylation of S707 and S708 is not conserved in other kinases and seems to be specific to PK3. The conserved residue T917, an analogue of T980 of mouse PERK that is present in the kinase activation loop and essential for eIF-2α phosphorylation, is of great interest (50). Unfortunately, there was little coverage of the protein in this domain in the two MS datasets, and we have no evidence of its phosphorylation. Notably, none of the single mutations we introduced completely abolished the auto- phosphorylation of PK3. However, the mutant PK3^F771L^ completely abolished the kinase activity.

As opposed to PERK, a transmembrane domain protein localized to the ER membrane, PK3 only associates with the ER, and upon activation, translocates to the nucleus. The translocation to the nucleus depends on its phosphorylation, since the translocation of a PK3 mutant defective in ATP binding was compromised. It is not currently known whether all the phosphorylation events on PK3 are auto- phosphorylations, or if other kinase(s) can act on PK3. Phosphorylation on PK3 by another kinase may still depend on recognition of auto-phosphorylated PK3, which does not occur in the PK3^F771L^ mutant.

A summary of the conditions or perturbations that induce SLS is presented in Fig. 7. This study provides evidence that SLS, which was initially discovered upon depletion of the SRP receptor, is activated by perturbation of a variety of factors involved in the translocation of proteins into the ER, their folding, and their modification along the intracellular protein trafficking pathway. In addition, SLS is also activated upon perturbations in the mitochondria. PK3 is therefore a key kinase in the cell, whose activation induces PCD leading to parasite elimination. PK3 should therefore be considered as an attractive drug target. A small molecule that would selectively activate PK3 could lead to suicide of these parasites.

**Figure 7.**
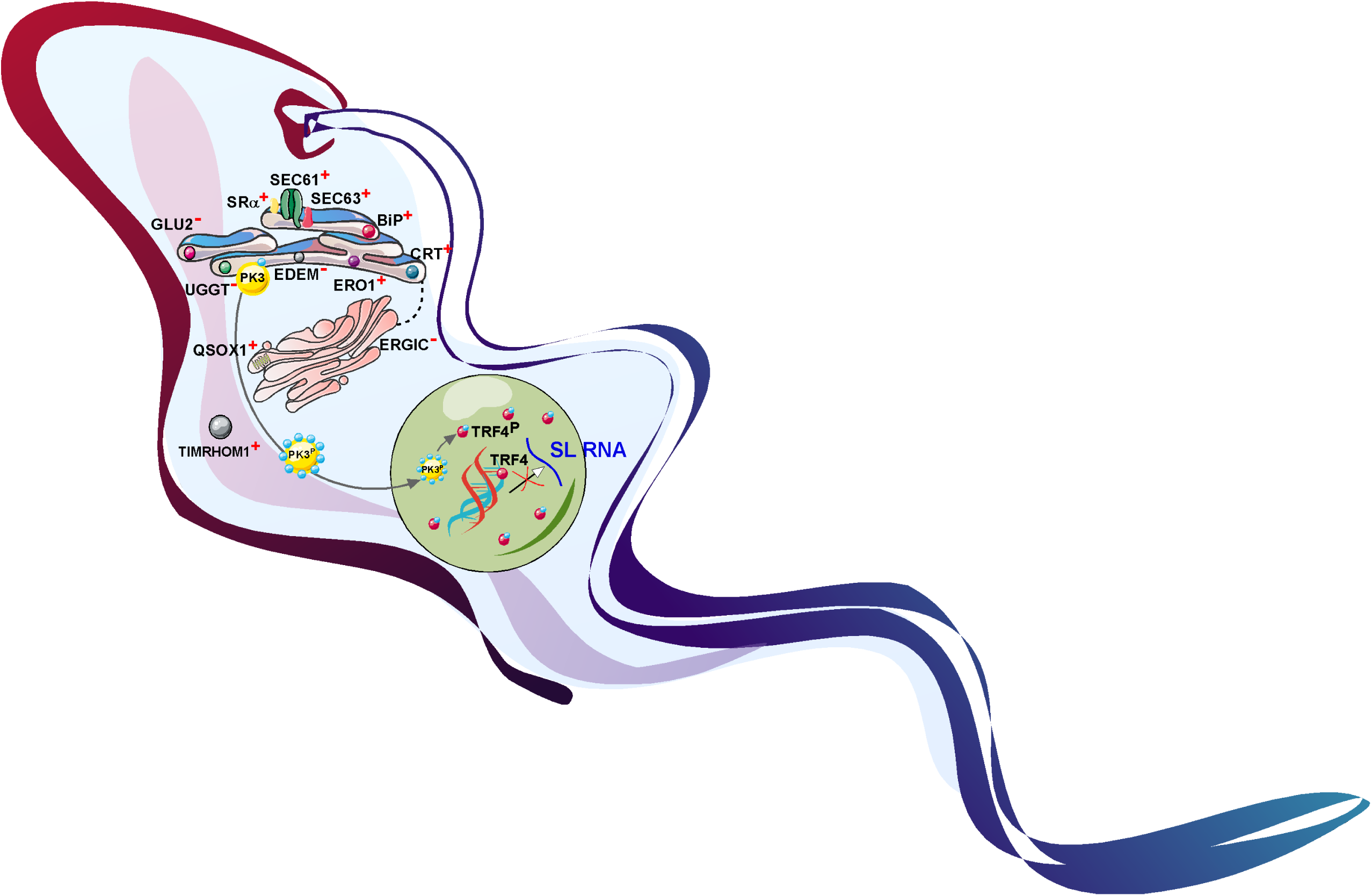
Protein factors whose depletion induces SLS and its mechanism. The scheme indicates the factors whose depletion induces SLS (marked with a plus (+) sign). Factors that do not induce SLS upon depletion are indicated with a minus (-) sign. During SLS, the PK3 undergoes phosphorylation on at least 9 sites. The PK3 migrates to the nucleus and phosphorylates TRF4, leading to detachment from its cognate promoter, and resulting in cessation of SL RNA transcription.

## ACKNOWLEDGMENTS

This study was supported by grants from the Israel Science Foundation to S.M. and D.F. S.M. holds the David and Inez Myers Chair in RNA silencing of diseases. Okalang Uthman is also affiliated to the Department of Biochemistry and Molecular Biology, Busitema University Faculty of Health Sciences, Mbale, Uganda.

## SUPPLEMENTARY LEGEND

**Table S1. List of primers used in this study.**

**Table S2. PK3 purification and Mass Spectrometry of the phosphorylated residues.** PK3-PTP was affinity purified as described in the Materials and Methods and in Figure 5. **A) Mass spectrometry analysis of proteins purified on titanium oxide column. B) Mass-spectrometry of isolated protein.** The protein was excised from a 10% SDS-polyacrylamide gel, and subjected to MS. The peptide sequences and phosphorylation sites obtained in **A** and **B** are indicated. PSMs: peptide spectrum matches, indicates the total number of identified peptide sequences. The Table S2 is provided as separate Microsoft Excel file.

**Figure S1.** Sequence alignment between *T. brucei* PK3, human PERK (PDB ID: 4G31) (56) and mouse PERK (PDB ID: 3QD2) (55) kinase domain. Removing the residues that are colored in black increases the sequence identity from 14.6% to 26.9%. Focusing on the ATP binding site (boxed in red) further increases the sequence identity to 63.6%. The conserved phenylalanine residues in the binding site of ATP are shown in a black box. Identical, similar and non-similar residues are colored in dark blue, light blue and white, respectively.

**Figure S2. Multiple sequence alignment PK3 to PERK kinase.** Amino acid sequences of human, mouse, *Drosophila*, *C. elegans* PERK and *T. brucei* PK1, 2 and 3 kinases were aligned using T-COFFEE Multiple Sequence Alignment Server (http://tcoffee.crg.cat/). The scores for the alignment are designated: BAD (indicated either in purple or green), AVERAGE (indicated in yellow) GOOD (indicated in pink). The domains are marked (I-V) as adopted from PROSITE (https://prosite.expasy.org/scanprosite/). The phosphorylated amino acid residues are indicated in red. The UniProt accession ID of HsPERK, MmPERK, DmPERK, CePERK, TbPK1, TbPK2 and TbPK3 are Q9NZJ5, Q7TQC8, Q9NIV1, Q19192, Q384V1, Q584D7 and Q583N6 respectively.

## REFERENCES

1. Michaeli S. 2011. Trans-splicing in trypanosomes: machinery and its impact on the parasite transcriptome. Future Microbiol 6:459–74.

2. Aphasizheva I, Aphasizhev R. 2016. U-Insertion/Deletion mRNA-Editing Holoenzyme: Definition in Sight. Trends Parasitol 32:144–156.

3. Clayton CE. 2016. Gene expression in Kinetoplastids. Curr Opin Microbiol 32:46–51.

4. Michaeli S. 2015. The response of trypanosomes and other eukaryotes to ER stress and the spliced leader RNA silencing (SLS) pathway in Trypanosoma brucei. Crit Rev Biochem Mol Biol 50:256–67.

5. Michaeli S. 2014. Non-coding RNA and the complex regulation of the trypanosome life cycle. Curr Opin Microbiol 20:146–152.

6. Michaeli S. 2012. Spliced leader RNA silencing (SLS) - a programmed cell death pathway in Trypanosoma brucei that is induced upon ER stress. Parasit Vectors 5:107.

7. Goldshmidt H, Matas D, Kabi A, Carmi S, Hope R, Michaeli S. 2010. Persistent ER stress induces the spliced leader RNA silencing pathway (SLS), leading to programmed cell death in Trypanosoma brucei. PLoS Pathog 6:e1000731.

8. Goldshmidt H, Sheiner L, Bütikofer P, Roditi I, Uliel S, Günzel M, Engstler M, Michaeli S. 2008. Role of protein translocation pathways across the endoplasmic reticulum in Trypanosoma brucei. J Biol Chem 283:32085–98.

9. Lustig Y, Sheiner L, Vagima Y, Goldshmidt H, Das A, Bellofatto V, Michaeli S. 2007. Spliced-leader RNA silencing: a novel stress-induced mechanism in Trypanosoma brucei. EMBO Rep 8:408–13.

10. Hope R, Ben-Mayor E, Friedman N, Voloshin K, Biswas D, Matas D, Drori Y, Günzl A, Michaeli S. 2014. Phosphorylation of the TATA-binding protein activates the spliced leader silencing pathway in Trypanosoma brucei. Sci Signal 7:ra85.

11. Hwang J, Qi L. 2018. Quality Control in the Endoplasmic Reticulum: Crosstalk between ERAD and UPR pathways. Trends Biochem Sci 43:593–605.

12. Michalak M, Groenendyk J, Szabo E, Gold LI, Opas M. 2009. Calreticulin, a multi-process calcium-buffering chaperone of the endoplasmic reticulum. Biochem J 417:651–66.

13. Pobre KFR, Poet GJ, Hendershot LM. 2019. The endoplasmic reticulum (ER) chaperone BiP is a master regulator of ER functions: Getting by with a little help from ERdj friends. J Biol Chem 294:2098–2108.

14. Field MC, Natesan SKA, Gabernet-Castello C, Koumandou VL. 2007. Intracellular trafficking in the trypanosomatids. Traffic 8:629–39.

15. Zito E. 2015. ERO1: A protein disulfide oxidase and H2O2 producer. Free Radic Biol Med 83:299–304.

16. McCaffrey K, Braakman I. 2016. Protein quality control at the endoplasmic reticulum. Essays Biochem 60:227–235.

17. Parodi AJ. 2000. Role of N-oligosaccharide endoplasmic reticulum processing reactions in glycoprotein folding and degradation. Biochem J 348 Pt 1:1–13.

18. Jones D, Mehlert A, Ferguson MAJ. 2004. The N-glycan glucosidase system in Trypanosoma brucei. Biochem Soc Trans 32:766–8.

19. Ito Y, Hagihara S, Matsuo I, Totani K. 2005. Structural approaches to the study of oligosaccharides in glycoprotein quality control. Curr Opin Struct Biol 15:481–9.

20. Kanehara K, Kawaguchi S, Ng DTW. 2007. The EDEM and Yos9p families of lectin-like ERAD factors. Semin Cell Dev Biol 18:743–50.

21. Wu X, Rapoport TA. 2018. Mechanistic insights into ER-associated protein degradation. Curr Opin Cell Biol 53:22–28.

22. Appenzeller-Herzog C, Hauri H-P. 2006. The ER-Golgi intermediate compartment (ERGIC): in search of its identity and function. J Cell Sci 119:2173–83.

23. Mehlert A, Zitzmann N, Richardson JM, Treumann A, Ferguson MA. 1998. The glycosylation of the variant surface glycoproteins and procyclic acidic repetitive proteins of Trypanosoma brucei. Mol Biochem Parasitol 91:145–52.

24. Manna PT, Boehm C, Leung KF, Natesan SK, Field MC. 2014. Life and times: synthesis, trafficking, and evolution of VSG. Trends Parasitol 30:251–8.

25. Tiengwe C, Muratore KA, Bangs JD. 2016. Surface proteins, ERAD and antigenic variation in Trypanosoma brucei. Cell Microbiol 18:1673–1688.

26. Field MC, Sergeenko T, Wang Y-N, Böhm S, Carrington M. 2010. Chaperone requirements for biosynthesis of the trypanosome variant surface glycoprotein. PLoS One 5:e8468.

27. Hope R, Egarmina K, Voloshin K, Waldman Ben-Asher H, Carmi S, Eliaz D, Drori Y, Michaeli S. 2016. Transcriptome and proteome analyses and the role of atypical calpain protein and autophagy in the spliced leader silencing pathway in Trypanosoma brucei. Mol Microbiol 102:1–21.

28. Harsman A, Oeljeklaus S, Wenger C, Huot JL, Warscheid B, Schneider A. 2016. The non-canonical mitochondrial inner membrane presequence translocase of trypanosomatids contains two essential rhomboid-like proteins. Nat Commun 7:13707.

29. Kornmann B, Currie E, Collins SR, Schuldiner M, Nunnari J, Weissman JS, Walter P. 2009. An ER-mitochondria tethering complex revealed by a synthetic biology screen. Science 325:477–81.

30. Szabadkai G, Bianchi K, Várnai P, De Stefani D, Wieckowski MR, Cavagna D, Nagy AI, Balla T, Rizzuto R. 2006. Chaperone-mediated coupling of endoplasmic reticulum and mitochondrial Ca2+ channels. J Cell Biol 175:901– 11.

31. Wang Z, Morris JC, Drew ME, Englund PT. 2000. Inhibition of Trypanosoma brucei gene expression by RNA interference using an integratable vector with opposing T7 promoters. J Biol Chem 275:40174–9.

32. Brun R, Schönenberger. 1979. Cultivation and in vitro cloning or procyclic culture forms of Trypanosoma brucei in a semi-defined medium. Short communication. Acta Trop 36:289–92.

33. Schimanski B, Nguyen TN, Günzl A. 2005. Highly efficient tandem affinity purification of trypanosome protein complexes based on a novel epitope combination. Eukaryot Cell 4:1942–50.

34. Fisk JC, Sayegh J, Zurita-Lopez C, Menon S, Presnyak V, Clarke SG, Read LK. 2009. A type III protein arginine methyltransferase from the protozoan parasite Trypanosoma brucei. J Biol Chem 284:11590–11600.

35. Alsford S, Horn D. 2008. Single-locus targeting constructs for reliable regulated RNAi and transgene expression in Trypanosoma brucei. Mol Biochem Parasitol 161:76–9.

36. Barth S, Hury A, Liang X, Michaeli S. 2005. Elucidating the Role of H/ACA-like RNAs in trans -Splicing and rRNA Processing via RNA Interference Silencing of the Trypanosoma brucei CBF5 Pseudouridine Synthase. J Biol Chem 280:34558–34568.

37. Liang X-H, Uliel S, Hury A, Barth S, Doniger T, Unger R, Michaeli S. 2005. A genome-wide analysis of C/D and H/ACA-like small nucleolar RNAs in Trypanosoma brucei reveals a trypanosome-specific pattern of rRNA modification. RNA 11:619–45.

38. He CY, Ho HH, Malsam J, Chalouni C, West CM, Ullu E, Toomre D, Warren G. 2004. Golgi duplication in Trypanosoma brucei. J Cell Biol 165:313–21.

39. Demmel L, Melak M, Kotisch H, Fendos J, Reipert S, Warren G. 2011. Differential selection of Golgi proteins by COPII Sec24 isoforms in procyclic Trypanosoma brucei. Traffic 12:1575–91.

40. Alon A, Grossman I, Gat Y, Kodali VK, DiMaio F, Mehlman T, Haran G, Baker D, Thorpe C, Fass D. 2012. The dynamic disulphide relay of quiescin sulphydryl oxidase. Nature 488:414–8.

41. Stern MZ, Gupta SK, Salmon-Divon M, Haham T, Barda O, Levi S, Wachtel C, Nilsen TW, Michaeli S. 2009. Multiple roles for polypyrimidine tract binding (PTB) proteins in trypanosome RNA metabolism. RNA 15:648–65.

42. Eliaz D, Kannan S, Shaked H, Arvatz G, Tkacz ID, Binder L, Waldman Ben-Asher H, Okalang U, Chikne V, Cohen-Chalamish S, Michaeli S. 2017. Exosome secretion affects social motility in Trypanosoma brucei. PLoS Pathog 13:e1006245.

43. Bangs JD, Uyetake L, Brickman MJ, Balber AE, Boothroyd JC. 1993. Molecular cloning and cellular localization of a BiP homologue in Trypanosoma brucei. Divergent ER retention signals in a lower eukaryote. J Cell Sci 105 (Pt 4:1101–13.

44. Bossard G, Grébaut P, Thévenon S, Séveno M, Berthier D, Holzmuller P. 2016. Cloning, expression, molecular characterization and preliminary studies on immunomodulating properties of recombinant Trypanosoma congolense calreticulin. Infect Genet Evol 45:320–331.

45. Palfi Z, Schimanski B, Günzl A, Lücke S, Bindereif A. 2005. U1 small nuclear RNP from Trypanosoma brucei: a minimal U1 snRNA with unusual protein components. Nucleic Acids Res 33:2493–503.

46. van Engeland M, Nieland LJ, Ramaekers FC, Schutte B, Reutelingsperger CP. 1998. Annexin V-affinity assay: a review on an apoptosis detection system based on phosphatidylserine exposure. Cytometry 31:1–9.

47. Ilani T, Alon A, Grossman I, Horowitz B, Kartvelishvily E, Cohen SR, Fass D. 2013. A secreted disulfide catalyst controls extracellular matrix composition and function. Science 341:74–6.

48. Demmel L, Melak M, Kotisch H, Fendos J, Reipert S, Warren G. 2011. Differential selection of Golgi proteins by COPII Sec24 isoforms in procyclic Trypanosoma brucei. Traffic 12:1575–91.

49. Ho HH, He CY, de Graffenried CL, Murrells LJ, Warren G. 2006. Ordered assembly of the duplicating Golgi in Trypanosoma brucei. Proc Natl Acad Sci U S A 103:7676–81.

50. Cui W, Li J, Ron D, Sha B. 2011. The structure of the PERK kinase domain suggests the mechanism for its activation. Acta Crystallogr D Biol Crystallogr 67:423–8.

51. Su Q, Wang S, Gao HQ, Kazemi S, Harding HP, Ron D, Koromilas AE. 2008. Modulation of the eukaryotic initiation factor 2 alpha-subunit kinase PERK by tyrosine phosphorylation. J Biol Chem 283:469–75.

52. Hornbeck P V, Kornhauser JM, Tkachev S, Zhang B, Skrzypek E, Murray B, Latham V, Sullivan M. 2012. PhosphoSitePlus: a comprehensive resource for investigating the structure and function of experimentally determined post-translational modifications in man and mouse. Nucleic Acids Res 40:D261–70.

53. Moraes MCS, Jesus TCL, Hashimoto NN, Dey M, Schwartz KJ, Alves VS, Avila CC, Bangs JD, Dever TE, Schenkman S, Castilho BA. 2007. Novel membrane-bound eIF2alpha kinase in the flagellar pocket of Trypanosoma brucei. Eukaryot Cell 6:1979–91.

54. Harding HP, Zhang Y, Ron D. 1999. Protein translation and folding are coupled by an endoplasmic-reticulum-resident kinase. Nature 397:271–4.

55. Cui W, Li J, Ron D, Sha B. 2011. The structure of the PERK kinase domain suggests the mechanism for its activation. Acta Crystallogr D Biol Crystallogr 67:423–8.

56. Axten JM, Medina JR, Feng Y, Shu A, Romeril SP, Grant SW, Li WHH, Heerding DA, Minthorn E, Mencken T, Atkins C, Liu Q, Rabindran S, Kumar R, Hong X, Goetz A, Stanley T, Taylor JD, Sigethy SD, Tomberlin GH, Hassell AM, Kahler KM, Shewchuk LM, Gampe RT. 2012. Discovery of 7- methyl-5-(1-{[3-(trifluoromethyl)phenyl]acetyl}-2,3-dihydro-1H-indol-5-yl)- 7H-pyrrolo[2,3-d]pyrimidin-4-amine (GSK2606414), a potent and selective first-in-class inhibitor of protein kinase R (PKR)-like endoplasmic reticulum kinase (PERK). J Med Chem 55:7193–207.

57. Webb B, Sali A. 2016. Comparative Protein Structure Modeling Using MODELLER. Curr Protoc Bioinforma 54:5.6.1–5.6.37.

58. Martí-Renom MA, Stuart AC, Fiser A, Sánchez R, Melo F, Sali A. 2000. Comparative protein structure modeling of genes and genomes. Annu Rev Biophys Biomol Struct 29:291–325.

59. Wiederstein M, Sippl MJ. 2007. ProSA-web: interactive web service for the recognition of errors in three-dimensional structures of proteins. Nucleic Acids Res 35:W407–10.

60. Friesner RA, Banks JL, Murphy RB, Halgren TA, Klicic JJ, Mainz DT, Repasky MP, Knoll EH, Shelley M, Perry JK, Shaw DE, Francis P, Shenkin PS. 2004. Glide: a new approach for rapid, accurate docking and scoring. 1. Method and assessment of docking accuracy. J Med Chem 47:1739–49.

61. Halgren TA, Murphy RB, Friesner RA, Beard HS, Frye LL, Pollard WT, Banks JL. 2004. Glide: a new approach for rapid, accurate docking and scoring. 2. Enrichment factors in database screening. J Med Chem 47:1750–9.

62. Mehnert M, Sommer T, Jarosch E. 2014. Der1 promotes movement of misfolded proteins through the endoplasmic reticulum membrane. Nat Cell Biol 16:77–86.

63. Lemberg MK. 2013. Sampling the membrane: function of rhomboid-family proteins. Trends Cell Biol 23:210–7.

64. Michel AH, Kornmann B. 2012. The ERMES complex and ER-mitochondria connections. Biochem Soc Trans 40:445–50.

65. Schnarwiler F, Niemann M, Doiron N, Harsman A, Käser S, Mani J, Chanfon A, Dewar CE, Oeljeklaus S, Jackson CB, Pusnik M, Schmidt O, Meisinger C, Hiller S, Warscheid B, Schnaufer AC, Ochsenreiter T, Schneider A. 2014. Trypanosomal TAC40 constitutes a novel subclass of mitochondrial β-barrel proteins specialized in mitochondrial genome inheritance. Proc Natl Acad Sci U S A 111:7624–9.

66. Ramakrishnan S, Asady B, Docampo R. 2018. Acidocalcisome-Mitochondrion Membrane Contact Sites in Trypanosoma brucei. Pathog (Basel, Switzerland) 7.

67. Rainbolt TK, Saunders JM, Wiseman RL. 2014. Stress-responsive regulation of mitochondria through the ER unfolded protein response. Trends Endocrinol Metab 25:528–37.

68. Lebeau J, Saunders JM, Moraes VWR, Madhavan A, Madrazo N, Anthony MC, Wiseman RL. 2018. The PERK Arm of the Unfolded Protein Response Regulates Mitochondrial Morphology during Acute Endoplasmic Reticulum Stress. Cell Rep 22:2827–2836.

69. Tiengwe C, Koeller CM, Bangs JD. 2018. Endoplasmic reticulum-associated degradation and disposal of misfolded GPI-anchored proteins in Trypanosoma brucei. Mol Biol Cell 29:2397–2409.

70. Tiengwe C, Brown AENA, Bangs JD. 2015. Unfolded Protein Response Pathways in Bloodstream-Form Trypanosoma brucei? Eukaryot Cell 14:1094– 101.

71. Eliaz D, Kannan S, Shaked H, Arvatz G, Tkacz ID, Binder L, Waldman Ben-Asher H, Okalang U, Chikne V, Cohen-Chalamish S, Michaeli S. 2017. Exosome secretion affects social motility in Trypanosoma brucei. PLOS Pathog 13:e1006245.

72. Liu L, Liang X, Uliel S, Unger R, Ullu E, Michaeli S. 2002. RNA interference of signal peptide-binding protein SRP54 elicits deleterious effects and protein sorting defects in trypanosomes. J Biol Chem 277:47348–57.

73. Goldshmidt H, Sheiner L, Bütikofer P, Roditi I, Uliel S, Günzel M, Engstler M, Michaeli S. 2008. Role of protein translocation pathways across the endoplasmic reticulum in Trypanosoma brucei. J Biol Chem 283:32085–98.

74. Hornbeck P V, Zhang B, Murray B, Kornhauser JM, Latham V, Skrzypek E. 2015. PhosphoSitePlus, 2014: mutations, PTMs and recalibrations. Nucleic Acids Res 43:D512–20.

75. Meares GP, Liu Y, Rajbhandari R, Qin H, Nozell SE, Mobley JA, Corbett JA, Benveniste EN. 2014. PERK-Dependent Activation of JAK1 and STAT3 Contributes to Endoplasmic Reticulum Stress-Induced Inflammation. Mol Cell Biol 34:3911–3925.

76. Sharma K, D’Souza RCJ, Tyanova S, Schaab C, Wiśniewski JR, Cox J, Mann M. 2014. Ultradeep Human Phosphoproteome Reveals a Distinct Regulatory Nature of Tyr and Ser/Thr-Based Signaling. Cell Rep 8:1583–1594.

77. Su Q, Wang S, Gao HQ, Kazemi S, Harding HP, Ron D, Koromilas AE. 2008. Modulation of the Eukaryotic Initiation Factor 2 α-Subunit Kinase PERK by Tyrosine Phosphorylation. J Biol Chem 283:469–475.

78. Vander Mierde D, Scheuner D, Quintens R, Patel R, Song B, Tsukamoto K, Beullens M, Kaufman RJ, Bollen M, Schuit FC. 2007. Glucose Activates a Protein Phosphatase-1-Mediated Signaling Pathway to Enhance Overall Translation in Pancreatic β-Cells. Endocrinology 148:609–617.

79. Wu X, Tian L, Li J, Zhang Y, Han V, Li Y, Xu X, Li H, Chen X, Chen J, Jin W, Xie Y, Han J, Zhong C-Q. 2012. Investigation of Receptor interacting protein (RIP3)-dependent Protein Phosphorylation by Quantitative Phosphoproteomics. Mol Cell Proteomics 11:1640–1651.

80. Trost M, Sauvageau M, Herault O, Deleris P, Pomies C, Chagraoui J, Mayotte N, Meloche S, Sauvageau G, Thibault P. 2012. Posttranslational regulation of self-renewal capacity: insights from proteome and phosphoproteome analyses of stem cell leukemia. Blood 120:e17–e27.

81. Mounir Z, Krishnamoorthy JL, Wang S, Papadopoulou B, Campbell S, Muller WJ, Hatzoglou M, Koromilas AE. 2011. Akt Determines Cell Fate Through Inhibition of the PERK-eIF2 Phosphorylation Pathway. Sci Signal 4:ra62–ra62.

82. Yamani L, Latreille M, Larose L. 2014. Interaction of Nck1 and PERK phosphorylated at Y 561 negatively modulates PERK activity and PERK regulation of pancreatic β-cell proinsulin content. Mol Biol Cell 25:702–711.

83. Urbaniak MD, Guther MLS, Ferguson MAJ. 2012. Comparative SILAC Proteomic Analysis of Trypanosoma brucei Bloodstream and Procyclic Lifecycle Stages. PLoS One 7:e36619.

84. Urbaniak MD, Martin DMA, Ferguson MAJ. 2013. Global quantitative SILAC phosphoproteomics reveals differential phosphorylation is widespread between the procyclic and bloodstream form lifecycle stages of Trypanosoma brucei. J Proteome Res 12:2233–44.

85. Mertins P, Qiao JW, Patel J, Udeshi ND, Clauser KR, Mani DR, Burgess MW, Gillette MA, Jaffe JD, Carr SA. 2013. Integrated proteomic analysis of post-translational modifications by serial enrichment. Nat Methods 10:634–637.

86. Shiromizu T, Adachi J, Watanabe S, Murakami T, Kuga T, Muraoka S, Tomonaga T. 2013. Identification of Missing Proteins in the neXtProt Database and Unregistered Phosphopeptides in the PhosphoSitePlus Database As Part of the Chromosome-Centric Human Proteome Project. J Proteome Res 12:2414– 2421.

87. Rigbolt KTG, Prokhorova TA, Akimov V, Henningsen J, Johansen PT, Kratchmarova I, Kassem M, Mann M, Olsen J V., Blagoev B. 2011. System-Wide Temporal Characterization of the Proteome and Phosphoproteome of Human Embryonic Stem Cell Differentiation. Sci Signal 4:rs3–rs3.

88. Hornbeck P V., Kornhauser JM, Tkachev S, Zhang B, Skrzypek E, Murray B, Latham V, Sullivan M. 2012. PhosphoSitePlus: a comprehensive resource for investigating the structure and function of experimentally determined post-translational modifications in man and mouse. Nucleic Acids Res 40:D261– D270.

89. Soderquist RS, Danilov A V., Eastman A. 2014. Gossypol Increases Expression of the Pro-apoptotic BH3-only Protein NOXA through a Novel Mechanism Involving Phospholipase A2, Cytoplasmic Calcium, and Endoplasmic Reticulum Stress. J Biol Chem 289:16190–16199.

90. Chen C-H, Shaikenov T, Peterson TR, Aimbetov R, Bissenbaev AK, Lee S-W, Wu J, Lin H-K, Sarbassov DD. 2011. ER Stress Inhibits mTORC2 and Akt Signaling Through GSK-3 -Mediated Phosphorylation of Rictor. Sci Signal 4:ra10–ra10.

91. Kebache S, Cardin E, Nguyên DT, Chevet E, Larose L. 2004. Nck-1 Antagonizes the Endoplasmic Reticulum Stress-induced Inhibition of Translation. J Biol Chem 279:9662–9671.

92. Krishnan N, Fu C, Pappin DJ, Tonks NK. 2011. H2S-Induced sulfhydration of the phosphatase PTP1B and its role in the endoplasmic reticulum stress response. Sci Signal 4:ra86.

